# Emergence of robust global modules from local interactions and smooth gradients

**DOI:** 10.1101/2021.10.28.466284

**Authors:** Mikail Khona, Sarthak Chandra, Ila Fiete

**Author notes:** Equal contribution (Mikail Khona, Sarthak Chandra).

## Abstract

Modular structure and function are ubiquitous in biology, from the scale of ecosystems to the organization of animal bodies and brains. However, the mechanisms of modularity emergence over development remain unclear. Here we introduce the principle of *peak selection*, a process in which two local interactions self-organize discontinuous module boundaries from a smooth global gradient, unifying the positional hypothesis and the Turing pattern formation hypothesis for morphogenesis. Applied to the brain’s grid cell networks, peak selection results in the spontaneous emergence of functionally distinct modules with discretely spaced spatial periods. Applied to ecological systems, a generalization of the process results in discrete systems-level niches. The dynamics exhibits emergent self-scaling to variations in system size and “topological robustness” [1] that renders module emergence and module properties insensitive to most parameters. Peak selection substantially ameliorates the fine-tuning requirement of continuous attractor dynamics even within single modules. It makes a detail-independent prediction that grid module period ratios should approximate adjacent integer ratios, furnishing the most accurate match to data to date, with additional predictions to connect physiology, connectomics, and transcriptomics data. In sum, our results indicate that local competitive interactions combined with low-information global gradients can lead to robust global module emergence.

## INTRODUCTION

Modular structures are ubiquitous in natural systems, from ecological niches to human communities, and from body structures to circuits in the brain. This is probably so because they are robust to localized perturbations [2, 3], faster to adapt if the world requires sparse or modular changes [4], and permit flexible, high-capacity computation through compositionality [5–10]. In this sense, modularity is the crux of biological organization.

The prevalence of modularity raises critical questions about its evolutionary and developmental origins: From an evolutionary perspective, modular solutions to a given problem form a vanishingly small subset of all possible solutions, raising the question of how and why they are selected. From the perspective of development, which we take here, the question is how modular structures form, and whether module features such as size, number, and boundary locations need to be genetically instructed or spontaneously emerge through unfolding physical processes such as symmetry breaking.

One hypothesis for the developmental emergence of structure is the positional information hypothesis espoused by Lewis Wolpert: Gene expression generates spatial morphogen concentration gradients, and different downstream genes become activated by thresholding the morphogen concentration [11, 12]. In line with this hypothesis, body segmentation in Drosophila [13], Fig. 1a-c, is controlled by a family of genes that become activated at different concentrations of maternally deposited Bicoid RNA. This gradient precedes and causally guides spatial bands of gene expression whose boundaries demarcate segment boundaries. A distinct hypothesis by Alan Turing is the idea of spontaneous pattern emergence from local competitive interactions, minimizing the use of detailed genetic information [12, 14]. Supporting this hypothesis is evidence of digit formation via pattern formation in hand morphogenesis [15]. A clear example of Turing pattern formation comes from the grid cell system in the medial entorhinal cortex (MEC) of mammalian brains, Fig. 1d-f. MEC neurons form a triangular grid-like pattern of activity as a function of space when animals navigate. Underlying these spatially periodic responses are periodic activity patterns in the cortex [16, 17]. Continuous attractor neural network (CAN) models of grid cells are based on Turing patterning [18] and make a number of stringent predictions that are supported by evidence from many studies [19–23]. However, it is unclear whether and how these CAN models for single grid cell modules translate to the formation of multiple discrete modules of grid cells with distinct periods [24], Fig. 1i.

**FIG. 1.**
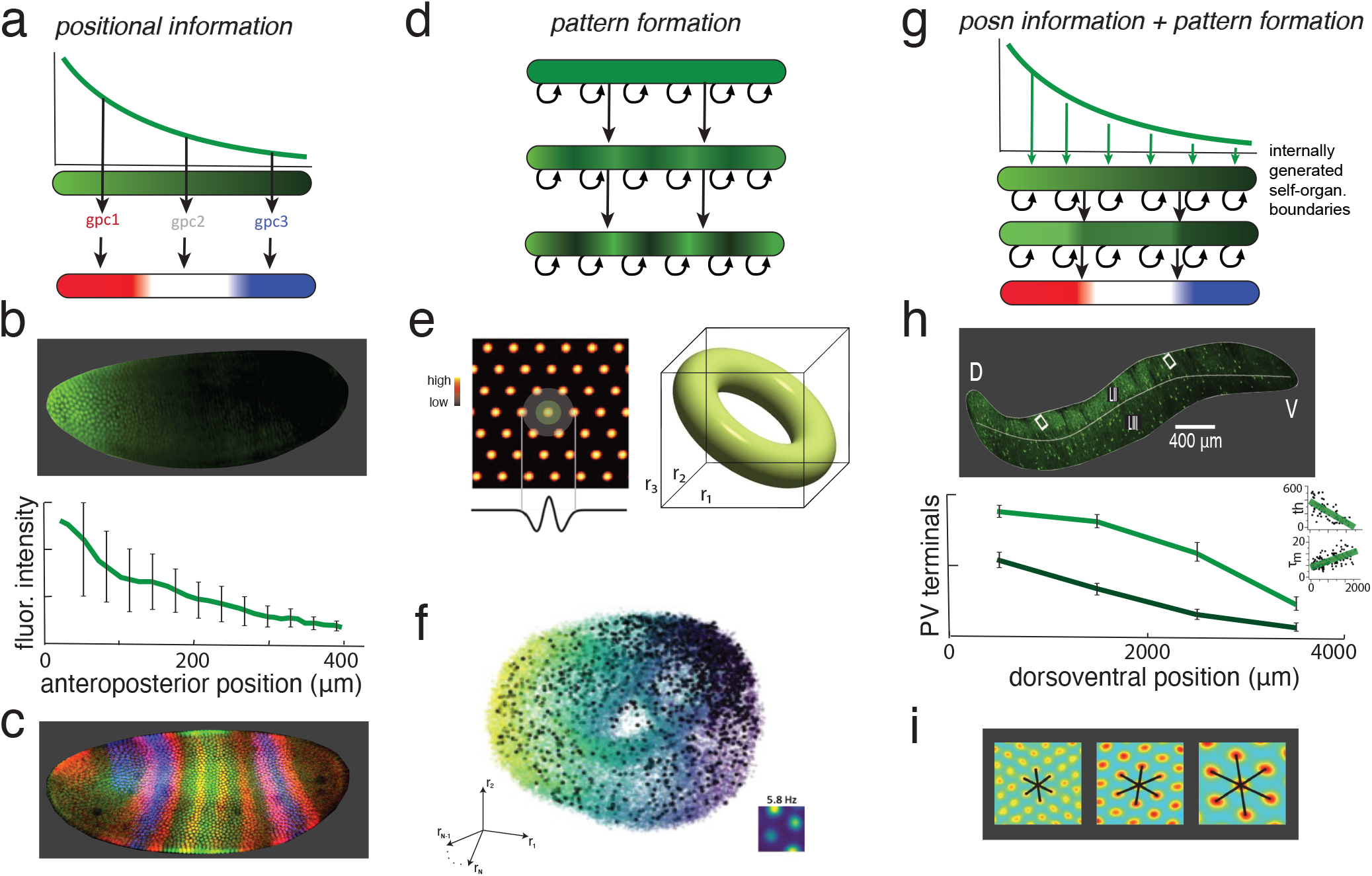
Positional versus pattern-forming mechanisms for structure formation and our hypothesis. (a) The positional hypothesis: global gradients are thresholded by different downstream gene expression cascades to generate structure [11]. (b) Fluorescence image of maternally deposited protein bcd RNA (based on maternal bicoid RNA deposition) early in development of the Drosophila embryo [46] sets up a polarity gradient. (c) A downstream gene-protein expression cascade, including gap and pair-rule genes, sets up body segment-defining bands by thresholding the bicoid gradient (immunofluorescence image adapted from [47]; segmentation figure adapted from [13].) (d) Spontaneous self-organized structure emergence (pattern formation) through competitive lateral interactions [14]. (e-f) The continuous attractor neural network (CAN) model for single grid cell modules [18] is based on Turing patterning based on local interactions and its predictions are consistent with the experimental data [19–21, 23]. These include the prediction of a continuous set of stable states with toroidal geometry across waking and sleep, and its recent confirmation [22]. (g) Our hypothesis: Positional and pattern forming mechanisms can interact to lead to structure emergence that exhibits the strengths of both mechanisms. (h-i) The long dorsoventral (DV) axis of medial entorhinal cortex (MEC; image of layers II and III) [32] exhibits smooth-seeming gradients in multiple cellular properties, while along the same axis, grid cells are organized into discrete modules with discontinuous jumps in their spatial periods (adapted from [24]).

The positional and Turing hypotheses individually have weaknesses [12, 25–27]: The positional mechanism is susceptible to noise in copy number [12, 28–30], and requires different downstream genetic cascades for each facet of structure that specify how and where that structure forms. Its implication that modular structure or function are driven by modularity in gene expression runs counter to at least some experimental studies that find continuous gradients in gene expression can underlie modular function [28, 31–44]. The pattern forming mechanism typically produces structure at a single scale given by the width of the local lateral interactions. These models do not possess scale invariance to (self-scaling with) the size of the system undergoing patterning.

Here, we hypothesize that positional and pattern forming mechanisms can be unified in a way such that modularity emerges through self-organization from local interactions without requiring modularity in gene expression, and the resulting process is scale-invariant, Fig. 1g. We show that such a process exists and is robust to most parametric variation and noise.

We focus first on the mammalian grid cell system, which exhibits structure on two scales: locally periodic patterns and global modules. The model produces strikingly accurate predictions about the sequence of successive spatial period ratios in grid cells, improving substantially on existing models. The process exhibits a “topological robustness” property that substantially eases the usual fine-tuning requirements of continuous attractor models of grid cells [45]. It also generates numerous predictions for future physiology, transcriptomics, and connectomics experiments.

Analyzing the underlying dynamical mechanisms of the process allows us to extract a general principle for global module emergence with smooth gradients and local interactions, which we call the *peak selection* principle. We then apply the peak selection principle to a very different problem, showing the emergence of modular multi-species niches in an interacting ecological system.

## GENERALIZATION OF CONTINUOUS ATTRACTOR MODELS FOR SINGLE GRID MODULES

Grid cells in the mammalian medial entorhinal cortex (MEC) of mammals exhibit spatially periodic response patterns as animals explore open spaces [16]. Moving ventrally along the long (dorsoventral or DV) axis of MEC, Fig. 1h, grid cells form modular and functionally independent subnetworks, with discrete jumps in spatial periodicity, Fig. 1i. Before considering mechanisms for the formation of multiple discrete grid modules, we extend the theory of single grid cell modules.

The dynamics of grid cells in a module are consistent with continuous attractor neural network (CAN) models [45]. CAN models involve pattern formation in neural activity through a linear (Turing) instability that is driven by strong local interactions between neurons. Realized predictions of CAN models include low-dimensional cell-cell relationships that are preserved across environments [19] and from waking to sleep [20, 21], as well as the direct demonstration of a toroidal set of stable states in the dynamics of single modules, Fig. 1e-f [22].

Existing CAN models show that grid tuning can arise two interaction profiles: a center excitation-surround inhibition (Mexican hat) shape (including an inhibition-only version) [18], or a uniform local inhibition shape (which we term a “Lincoln hat”) [48]. We derive a set of conditions that contain an infinte set of kernel shapes (SI A), which we hypothesize are sufficient for grid-like patterning. The conditions are simple constraints on the local interaction kernel *W* : that it is radially symmetric, that its integral is negative (inhibition dominated: *W* (*x, x*′)*dx*′ < 0), and that it is strong enough such that that there is some value of *k* > 0 such that its Fourier transform satisfies 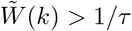. The latter can almost always be made to hold for interactions that satisfy the former by scaling *W* (SI Sec. A). This result significantly expands the generality of CAN models for single grid cell modules. Sampling randomly from interaction profiles satisfying these conditions and running the network dynamics with rectified linear neurons, we find that all sampled profiles produce grid-like patterning (Fig. 2a).

**FIG. 2.**
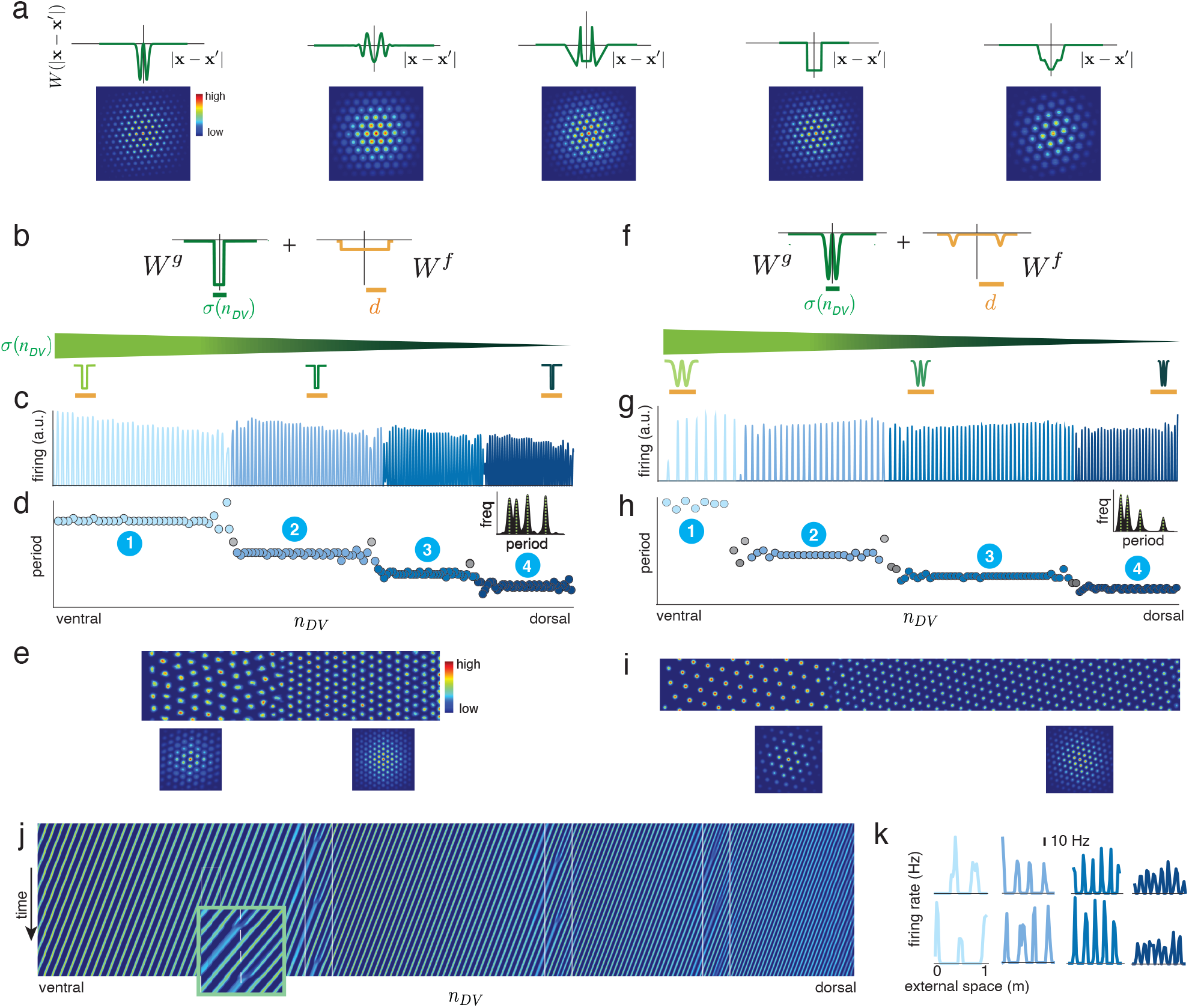
Two *local* interactions, with graded and fixed widths, respectively, lead to global module emergence. (a) Generalization of CAN grid cell models: 5 examples from an infinite set of distinct local interaction kernel shapes that can lead to grid-like patterning. (b) Combining two local interactions, one whose width (*σ*(*n*_*DV*_)) scales smoothly along the DV axis (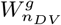, green) and a broader but still-local one whose width (*d*) remains fixed along the neural strip. Interaction widths indicated below the gradient are drawn to scale relative to the activity shown in (c). (c-e) The two interactions from (b) lead to spontaneous emergence of modules with distinct periods in 1-dimensional (c) neural strip, with extracted periods shown in (d). The same kernels applied to a 2-dimensional neural sheet (e), with the 2d autocorrelation function of the local (single-module) patterns in the neural sheet (bottom). (f-i) Same as (c-e), but for a different pair of interaction kernels 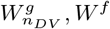 with distinct gradient shape *σ*(*n*_*DV*_) and endpoints (*σ*_min_, *σ*_max_) from (c-e). (j) The response of the 1-dimensional neural strip shown over time when the network is driven by a smoothly graded velocity input, white lines highlighting the temporal evolution of dynamics at the module boundary (inset: magnification of the first boundary). (k) The independent velocity-driven pattern dynamics in each module result in regular periodic spatial tuning curves (shown are 2 cells per module). *See Methods for parameter and simulation details*.

## FIXED AND GRADED LOCAL INTERACTIONS FOR GLOBAL MODULARITY EMERGENCE

Physiological experiments reveal biophysical gradients along the DV axis[32, 38, 42], Fig. 1h, raising the possibility that combining such a gradient with CAN pattern formation mechanisms might lead to the emergence of multiple discrete modules. We replaced the translation-invariant interaction kernels in CAN grid cell models by kernels with a slowly increasing width *σ*(*n*_*DV*_) along the DV axis of the model neural sheet, SI Fig.11a, where *n*_*DV*_ refers to DV location (while **x** refers to the general 2-dimensional position on the neural strip):

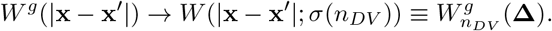

In numerical simulations, this slow DV variation of the local interaction width in the CAN model produces hexagonally arranged activity bumps with a growing period (SI Fig. 11b). However, the variation in pattern period is smooth, reflecting the smooth width gradient: there is no emergent modularization (SI Fig. 11c).

Given that global modularity and local patterning together cover two spatial scales, it might be necessary to include two scales in the lateral interactions. However, just as local interactions lead to globally periodic structure, we hypothesize that adding a second but still *local* interaction might be sufficient to induce globally modular structure.

The meaning of a second interaction must be distinct from merely a shape difference relative to the first, because the sum of two local kernels is otherwise just another local kernel, and the graded period result of SI Fig. 11b-c will still hold. Therefore, we explored adding a second type of local lateral interaction, *W*^*f*^, which remains fixed across the DV axis while the first is graded. The combined interaction is:

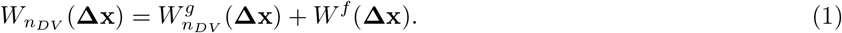

We assume that the fixed interaction is wider than the graded interaction, but both interactions are local and much smaller than the length of the cortical sheet: *σ*_max_ ≤ *d* ≪ *L*, where *d* is the width of *W*^*f*^ and *L* is the DV length of the neural sheet. Remarkably, the addition of such a fixed-scale interaction leads the network to spontaneously decompose into a few discrete modules, with coherent periodic activity patterns locally and discontinuous jumps in period globally, Fig. 2b-d. As with single module grid patterning, there is broad latitude in the shapes of the interaction kernels, *W*^*g*^ and *W*^*f*^, so long as one is graded and the other is wider but fixed in width along the DV axis, Fig. 2f-h. The same combination of a graded-width and a fixed-scale interaction produces robust and spontaneous decomposition of dynamics into discrete modules across 1D and 2D network models, Fig. 2e,i. (See SI Sec. D8 for additional model results for 2D networks.)

The modules that emerge are much larger than both local interaction scales, cf. Fig. 2b, f (where the interaction widths are rendered to scale), versus the formed modules in Fig. 2c-e, g-i. Given this surprising result of an emergent scale that is not directly related to the underlying physical interaction scales, we seek its causes below.

### Formed modules are functionally independent

The emergent modules, which are distinct in the periodicity of their patterning, are also functionally independent units. In single-module CAN models, providing cells with a velocity input causes the pattern to flow in a direction and with a speed proportional to the velocity input — the network can thus track current location by integrating a velocity signal [49–52], as grid cells are believed to do [18]. For the distinct modules to independently perform velocity integration means that their patterns flow independently, even though connectivity is as continuous across module boundaries as it is within modules, which seems intuitively implausible. However, when we drive all modules with a common velocity input, we find that the patterns in each module flow independently. The boundaries between modules remain sharp and fixed at the same cortical locations, with a spatially discontinuous phase dislocation between boundaries with module phases separately updating and shifting relative to each other in time, Fig. 2j. This dynamics results in veridical, independent velocity integration within and across modules, so that all cells (even those close to the module boundaries) have periodic spatial tuning curves, Fig. 2k.

## ANALYTICAL THEORY OF MODULARIZATON: PEAK SELECTION, TOPOLOGICAL ROBUSTNESS, AND SELF-SCALING

The generality and robustness with which discrete modules emerge from the combination of a fixed-scale and a graded-scale local interaction — with very different kernel shapes as in Fig.2b,f — suggests a general principle at work. Starting from an initial condition of uniform activity, the network exhibits nearly immediate (within 1-2 biophysical time-constants *τ*) signs of modularization, Fig. 3a. Modularization begins before most neurons have crossed their nonlinear thresholds, and unfolds concurrently with local periodic patterning (Fig. 3a,e). The system also exhibits localized eigenvectors (SI sec G; similar to the Anderson localization phase transition in condensed matter physics [53]). Both phenomena point to a linear instability-driven modularization mechanism, hinting at the possibility of a unified theory for patterning and modularity. We derive such a theory, summarizing it below with details in SI (Sec.B). Besides establishing how, why, and when modularity emerges, the theory accurately predicts the discrete pattern periods of all modules, the number and sizes of modules, and the locations of module boundaries (explored below).

**FIG. 3.**
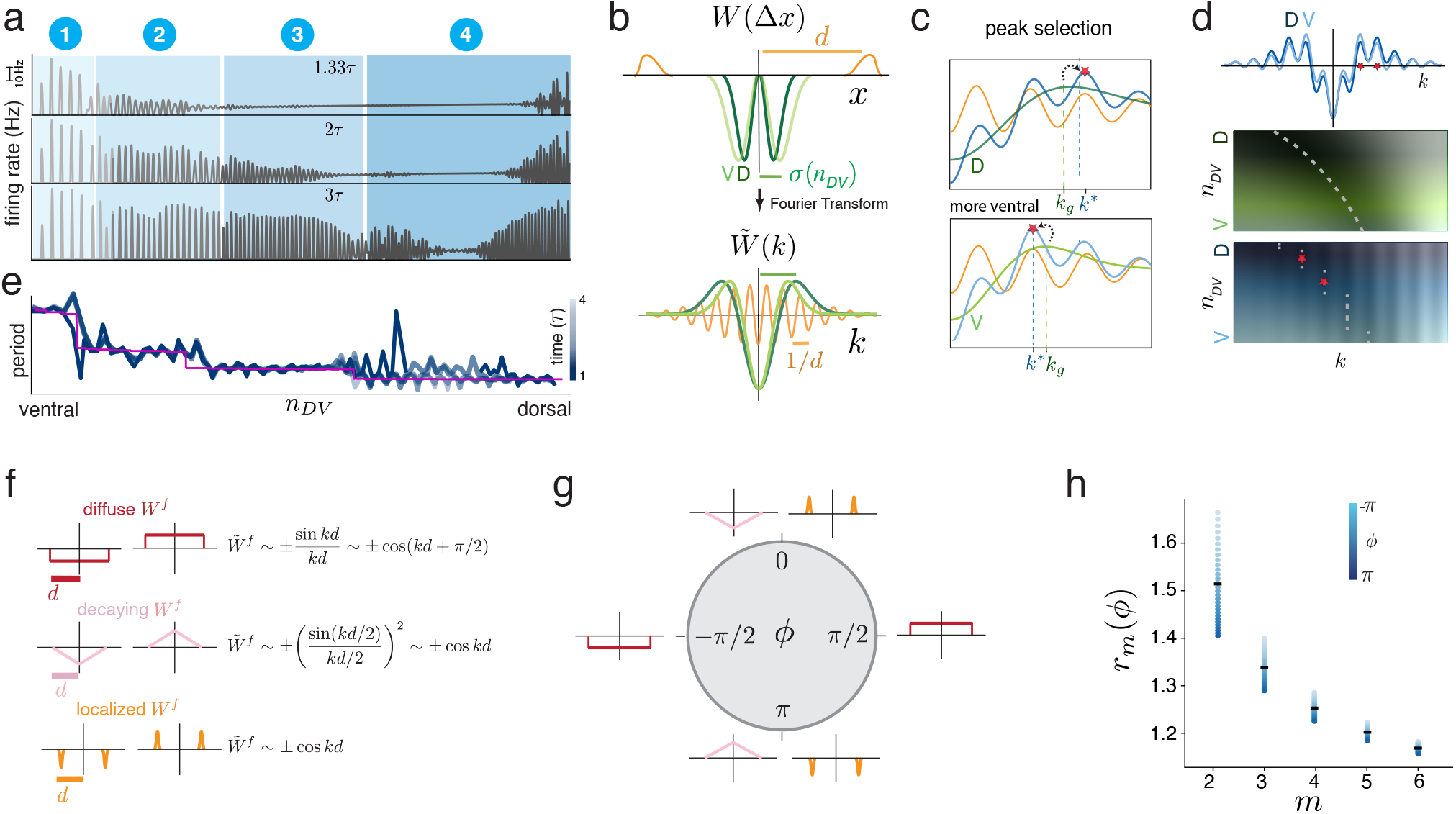
Theory of module emergence: multi-scale linear instability and topological peak selection. (a) Snapshots of population activity within a few neural time-constants (*τ*) of initializing the dynamics at a uniform state. Modules appear in situ at the same time as local patterning, before most neurons have hit their nonlinear thresholds. (b) Top: Schematics of fixed-width (orange) and graded (green) interaction kernels. Kernels from two different DV locations (designated D for dorsal and V for ventral). Bottom: their Fourier transforms. (c) Peak selection process: The global maxima in Fourier space (blue) are based on combining the graded interaction (green) with the fixed interaction (orange). As the green peak slides across, the global maximum (marked by red star) jumps abruptly from the position of one orange peak to the next. (d) Top: Summed Fourier transform of the two local interactions (darker (lighter) blue: more dorsal (ventral)). Middle: The location of the maximum of the graded interaction varies smoothly as a function of DV location. Bottom: the maximum of the summed interaction jumps discontinuously (bottom). (e) Dark to light blue curves: activity pattern periods from (a) for early to late times after initialization. Module boundaries and periods remain unchanged from the earliest time-points. Pink: theoretical prediction of periods and module boundaries from Eq. 2. (f) Left: Example simple fixed-scale interaction profiles that produce modularization: profiles can be roughly categorized as diffuse, decaying, or localized. Right: the dominant terms in their Fourier transforms. (g) The Fourier phases of the interactions in (a). (h) Theoretically predicted sequence of period ratios for any value of *ϕ* (blue circles), for module numbers 2-6. Any dependence on *ϕ* and thus the shape of the fixed-scale interaction is weaker for higher module numbers (smaller period/dorsal modules). *See Methods for parameter and simulation details for* (*a*) *and* (*e*).

The theory examines how infinitesimal perturbations evolve from an unstable initial fixed point *s*_0_(*n*_*DV*_). In spatially local windows that are much larger than the local interaction scale *σ*(*n*_*DV*_),*d* but much smaller than the full system size *L*, the dynamics can be decomposed into local Fourier modes (SI sectionD 8) and solved because the spatial variation in the interaction 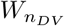 of Eq. (1) is slow. Under the same simple conditions on the interaction kernels given above (in Generalization of continuous attractor models for single grid cell modules), the network forms a *spatially varying* patterned state with the following inverse periods (wave vectors) as a function of the DV location *n*_*DV*_ :

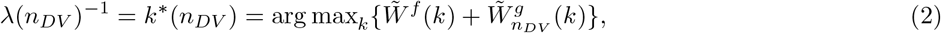

where 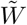 denotes the Fourier transform of *W*.

The spatial structure of this patterned solution can be gleaned from the interplay between the two terms. Suppose the interaction kernel *W*^*f*^ is *simple*: primarily characterized by one length-scale *d* > *σ*_max_ and such that other length-scales are not prominent and much smaller than *d* (Fig. 3b, top, orange). Then (except for a set of hypothesized measure zero in the space of functions *W*^*f*^ ; SI Sec. Dfor details), its Fourier transform has an approximate form (∼cos(*kd* −*ϕ*)), with closely spaced peaks every ∼ 1/*d* (Fig. 3b, bottom, orange) and a phase *ϕ*. These local maxima remain the same across the neural strip because *W*^*f*^ is not graded.

By contrast, 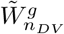 exhibits a broad Fourier peak (of scale ∼ 1/*σ*(*n*_*DV*_) ≫ 1/*d*). The width and location of this peak smoothly contracts along the long axis (as *σ*(*n*_*DV*_) increases; Fig. 3b, bottom, green). This interaction drives spatial patterning and its graded variation is ultimately responsible for changes in period. However, the narrow peaks of 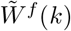 determine the set of potential locations of the global maximum, while the smoothly moving peak from 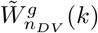 performs “peak selection” on these possibilities (Fig. 3c and SI Movie 1). It sweeps through the narrow local maxima of 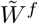, and as it does so one of the narrowly spaced peaks becomes the global maximum. As it sweeps on, the global maximum abruptly and discontinuously jumps to the next peak of 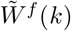, defining the abrupt changes in period between modules, Fig. 3c-d. Thus the spatial periods along the DV axis form a discrete set determined by the maxima of 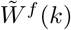, which occur at

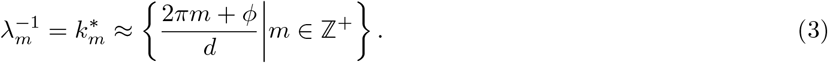

The module periods are proportional to the width *d* of the fixed-scale interaction *W*^*f*^, and are given by this scale divided by a set of integers *m*.

Permitted values of the integers *m* are determined by the set of local maxima *k*\*(*n*_*DV*_) of the fixed-scale interaction (in Eq. 2) that fall within the range [*η*/*σ*_max_, *η*/*σ*_min_], where *η* is a proportionality constant (SI Sec. D7). It follows that the number of allowed periods (and thus modules) is determined by the set of integers *m* that fit in the following interval:

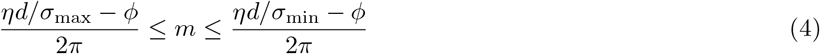

The phase *ϕ ϵ* [−*π, π*] is a simple scalar, determined by coarse-grained features of *W*^*f*^ (Fig. 3f-g): A diffuse *W*^*f*^ out to scale *d* leads to *ϕ* ≈ *±π*/2 for inhibitory or excitatory interactions respectively; a localized *W*^*f*^ at scale *d* leads to a value of *ϕ* close to 0 or *π* for excitatory or inhibitory interactions respectively; a decaying *W*^*f*^ also leads to *ϕ* close to 0 or *π* for inhibitory or excitatory interactions, respectively. Intermediate values of *ϕ* can be obtained by interpolating between these fixed-scale lateral interaction shapes (See Fig. 12 for examples of numerical simulations for several cases).

The analytical expression for module periods obtained by evaluating Eq. 2 on the Fourier transform of *W* exactly predicts the values from numerical simulation (Figs. 3e, 4b, SI Fig. 12). Similarly, the even simpler analytical expression for period in Eq. with *ϕ* computed from *W*^*f*^ and without free parameters, also exactly predicts module periods from numerical simulation across diverse lateral interaction shapes (Figs. 3a, 4b, SI Fig. 12).

### Period ratio prediction and parameter invariance through *topological protection*

The (inverse) module period expression of Eq. 3 supplies a quantitative prediction about module period ratios, which have been characterized experimentally [24] and are the subject of several theoretical models [54, 55]. In contrast to results in which period ratios have been described with one value regardless of the module number, our prediction varies with module: the period ratio of the *m*th module to the *m* + 1th module is given by:

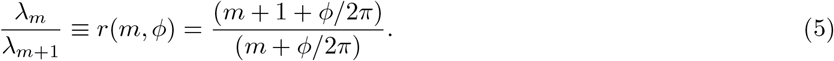

Note that while the module periods are determined by the scale *d* of the fixed-scale interaction, the module period ratios are strikingly and completely independent of any scale: neither *d*, 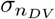, nor *L*. For *ϕ* = 0, The ratios of adjacent modules are simply successive integer ratios, with the integer indexing the module number (Fig. 3h). It is also independent of the functional form of gradation in 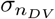.

The only parameter dependence in the period ratios is through the scalar phase *ϕ*, which itself depends only on coarse properties of the lateral interaction *W*^*f*^ (Fig. 3f-h). This extreme invariance of the period ratios to almost all parameters is a signature of a topologically protected[1] process (SI Sec.D 7), and is due to the discrete topology of the set of successive integers which are generated by the peak selection process. Thus, the modularization process is robust to nearly all parameteric variation.

### Emergent self-scaling of modules

Above we saw that within-module properties like pattern periods within each module are independent of details of the lateral interaction kernel shapes, the size *L* of the neural sheet, the width of the graded interaction and the functional form of the gradient. However, total independence of structure emergence from system parameters can be a weaknesses of pattern forming dynamics, if structures do not scale with quantities like system size [12, 56].

Interestingly and surprisingly, if the minimum and maximum widths (*σ*(0) = *σ*_max_, *σ*(*L*) = *σ*_min_) of the graded interaction remain fixed at the original absolute sizes, as does the absolute width of the fixed-scale interaction, while the network size *L* is varied, the number of formed modules remains unchanged, meaning that the module formation process has self-scaled to the system size (Fig.4a-b). Each module’s internal periodicity remains unchanged (Fig.4a-b), but the modules are now larger in width so that they each occupy the same fraction of the length of the neural sheet (Fig.4b). In other words, the dynamics of module formation exhibit an emergent invariance or self-scaling property with brain size, automatically adjusting to the size of the substrate without requiring any change in the biological and genetic parameters that control connectivity width.

**FIG. 4.**
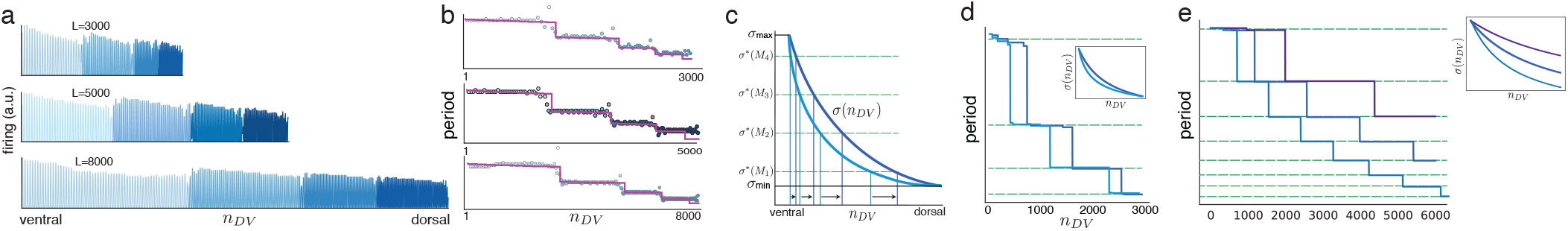
Emergent self-scaling. (a) Increasing the size of the neural sheet while holding constant the minimum and maximum graded interaction widths and the fixed interaction width, the within-module periods remain the same but module sizes expand so that the system has the same number of modules regardless of system size. (b) Extracted periods from results in (a). The neural axis is scaled (normalized) by network size to compare relative module sizes; the period axis is the same across plots (preserved periods in each module). Pink: analytical predictions from Eq. 2. (c) Different functions (shapes) for the monotonically graded interaction width *σ*(*n*_*DV*_) are predicted theoretically to result in the same number of modules if the minimum and maximum values of the width (*σ*_min_, *σ*_max_) remain unchanged. Shape changes only affect the detailed positions of module boundaries. (d) *k*\* calculated from numerical Fourier transform of interaction matrix with two different gradient shapes, holding *σ*_min_ and *σ*_max_ fixed. Module number and periods remain unchanged, while boundaries shift. (inset) The shapes of the gradient in the width of the primary pattern-forming interaction for the two choices of gradient shapes. Green dashed lines are scales corresponding to each local maxima of the secondary interaction. (e) *k*\* calculated from numerical Fourier transform of interaction matrix with three significantly different values of *σ*_min_ while holding the spatial extent of the system fixed. The number of formed modules changes from 3 to 5 to 8, while the periods of the first few modules (that are common across all three simulations) remain unchanged. (inset) The shapes of the gradient *σ*(*x*) in the primary pattern-forming interaction for the three choices of gradient shapes. *See Methods and SI SecD 7 for parameter and simulation details*.

We can understand this analytically within our theoretical framework: The number of formed modules *N*_*mod*_ is given by the number of integers satisfying Eq. (4), and thus is given by:

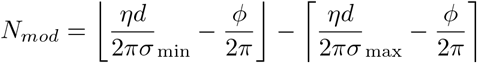

where ⌊, ⌋ ⌈ ⌉ indicate the floor and ceiling operations, respectively. In words, the number of modules is determined by the (integer) difference in the ratios of *d* with *σ*_min_, *σ*_max_: it is based on the interplay (difference) between the width ratios of the two local kernels, without depending on their specific widths or even the values of their ratios. Consistent with the numerical simulation results (Fig.4a-b), the number of formed modules is predicted to independent of the size of *L* if *σ*_min_, *σ*_max_,*d* are held fixed, which gives the dynamics its self-scaling property.

Even more so than for the within-module periods (cf. Eq. 3), the number of modules depends on the shapes of *W*^*f*^, *W*^*g*^ only slightly, through the scalar factor *ϕ*. Large variations in the shape result in the same number of modules (Fig. 2 and additional profiles in SI Fig.12). The number of modules is also notably predicted to not depend on the shape of the monotonically varying width function *σ*(*n*_*DV*_) (Fig. 4c).

We can make a more approximate but still fairly accurate prediction for where module boundaries will form (Fig. 3e, pink, and SI D 7):

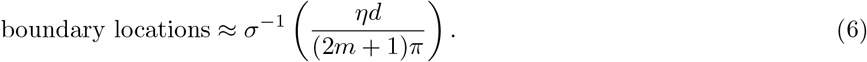

where *σ*^−1^ is the inverse function of *σ*(*n*_*DV*_), such that *σ*^−1^ ∘ *σ*(*n*_*DV*_) = *n*_*DV*_. Unlike the module number and period, the module boundary locations are predicted to depend on the shape of the gradient *σ*(*n*_*DV*_) (Fig. 3c). Numerical simulations confirm the independence of module number and period to changes in the shape of *σ*(*n*_*DV*_) when *σ*_min_, *σ*_max_ are fixed, but that boundary locations and therefore relative module sizes can change (Fig. 4d), see SI Sec D 7 for more details.

Finally, even if *σ*_min_, *σ*_max_ are varied smoothly (while *d* is held fixed), or if *d* is varied smoothly (while *σ*_min_, *σ*_max_ are held fixed), the number of modules will remain fixed until the change becomes large enough to accommodate one additional or one less module. All these features are properties of the topologically robust process of module emergence, and the number of modules is a topological invariant of the system [1].

A corollary of the independence of module number to system size is that the average module size will scale in proportion to system size, as *L*/*N*_*mod*_. Thus, if the neural sheet is large, the module sizes can be orders of magnitude larger than any of the lateral interaction scales, *σ*_min_, *σ*_max_, *d*, resolving the mystery of what sets the scale of individual modules, why they are unrelated to the local interaction scales, and why they are global in size, Fig. 2b-d,f-h.

Self-scaling occurs when holding the endpoints of the global gradient fixed as system size is varied, and is a property of positional information models. Thus, similar results could have been obtained by applying a set of thresholds to the function 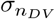, which then activated different downstream process to specify the boundaries and period for the corresponding module, which would correspond to the positional or french-flag model [11]. The difference here is that the modules are *self-generated* and do not require externally imposed thresholds or control. The self-organization process allows for mathematical prediction of the periods. An additional consequence for evolutionary dynamics is that changing the number of modules in the peak selection mechanism involves simply changing the value of one endpoint (*σ*_max_ or *σ*_min_) of the graded interaction function *σ*(*n*_*DV*_), rather than creating new gene expression cascades for each added module (Fig. 4e).

The simple analytical expressions for module period and period ratios, their parameter (in)dependence, and self-scaling properties accurately match the results of full numerical simulations, Fig. 5a.

**FIG. 5.**
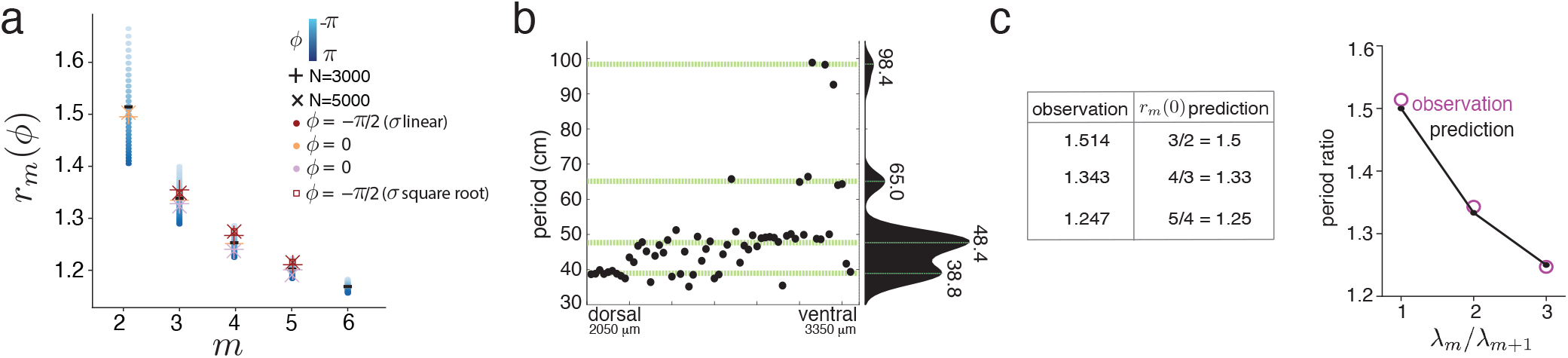
Comparison of precise period ratio predictions with data. (a) Period ratio predictions from 3h together with numerical simulation (other symbols) of neural circuit models with the set of fixed-scale interaction profiles shown in 3f, same color code. Numerical simulations with all combinations of network size and weight profiles are shown. (b) Observed periods of grid cells from multiple modules [24] (c) Successive period ratios computed from the observation (left column), and predicted period ratios for *ϕ* = 0 (middle column). Ratios match predicted values with *R*^2^ = 0.999 (right column). *See Methods for parameter and simulation details*.

### Neural data matches detailed predicted sequence of period ratios

We compare our predictions to existing results, which estimate a single ratio across adjacent module pairs [24], by averaging the predicted ratios across 4 modules and across different phases *ϕ* (SI Sec. D6). This yields a predicted value of 1.37, in good agreement with experimental results of an average ratio of ∼ 1.42 across animals as reported in[24] and 1.368 for the animal shown in Fig. 5b.

Further, we compare our more fine-grained successive period ratio predictions with published per-module period values. Our prediction with *ϕ* = 0 matches the sequence of observed period ratios [24] strikingly well, Fig.5c, also for other datasets in which multiple grid periods ratios are available from single individuals (SI Sec. D10).

## PEAK SELECTION ENHANCES ROBUSTNESS WITHIN INDIVIDUAL GRID MODULE ATTRACTOR NETWORKS

We have seen that peak selection-based emergence leads to module formation via patterning on two scales, and that the results are invariant to variations in parameters, function shapes, and the form of the global gradients. We next investigate and discover that the two-scale interaction and topological structure emergence process also confers substantial additional robustness to traditional continuous attractor models. Specifically, this mechanism makes continuous attractor dynamics resistant to several forms of weight heterogeneity and activity perturbation.

### Robustness within individual continuous attractor networks

Standard attractor models [18, 23, 57, 58] require weight homogeneity to generate regular patterns and continuous attractor states for representing continuous variables. Heterogeneities degrade the pattern and the continuity of attractor states. If the heterogeneities are sufficiently large [59, 60], pattern formation can entirely fail (Fig. 6a, left and SI Sec. D9 for details). This susceptibility of continuous attractor dynamics to noise is one of the fundamental open problems for continuous attractor models [23].

**FIG. 6.**
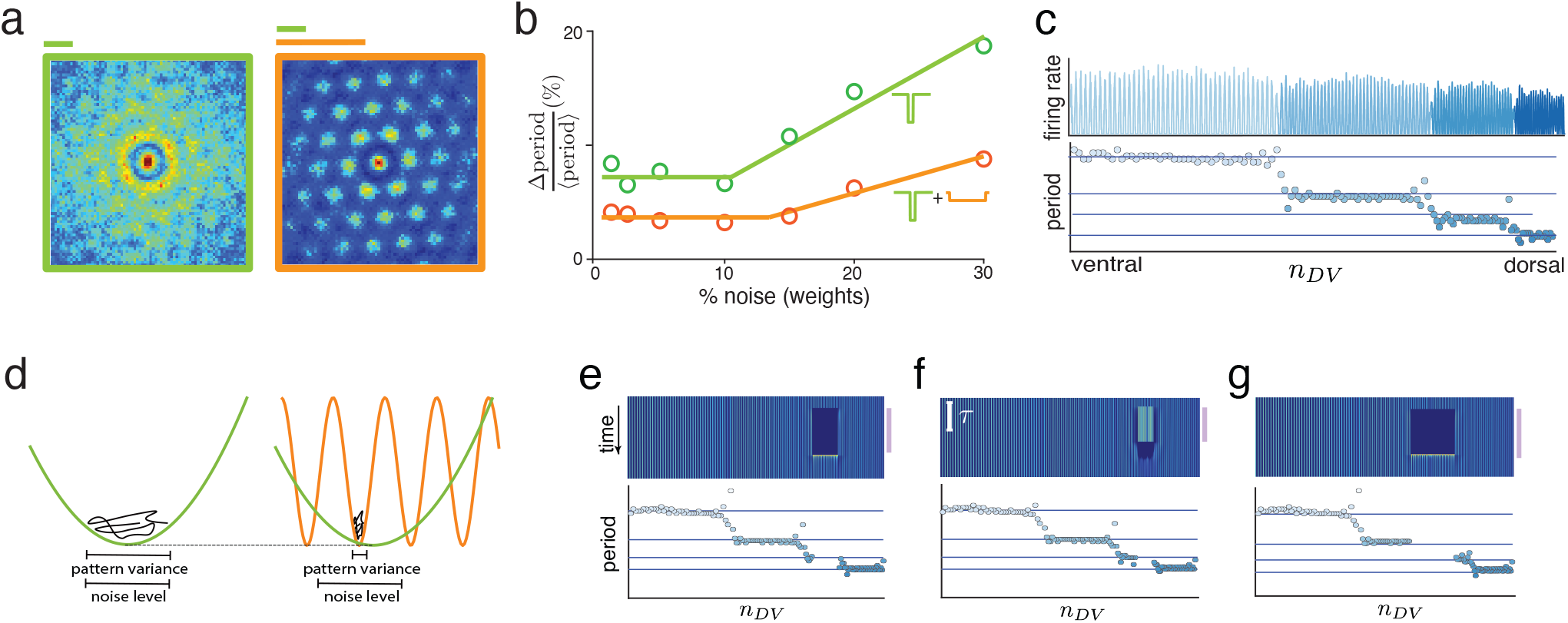
Enhanced robustness to weight heterogeneity, noise, and activity perturbation by peak selection. (a) Left: Weight heterogeneity (here, radial asymmetry and i.i.d. noise) quickly destroys discernable pattern structure in standard continuous attractor models [18]). (Simulation of a 100×100 neuron network with kernel *W*_*g*_ as in Fig. 2d-e; scale shown by green bar.) Right: Addition of a secondary wider local interaction (scale shown by orange bar), with noise in both sets of weights, rescues patterning. (b) Variability in 1-dimensional patterning versus the magnitude of added noise in the weights, for single-scale weights (green), and for networks with two local interaction scales (orange). Pattern variation is the ratio of the standard deviation to the mean of the pattern period. (c) The same as (a), showing regularity in period despite the addition of noise in a 1-dimensional setting. (d) The mechanism for enhanced within-module robustness: the broader local interaction scale enforces a narrower set of solutions in the energy landscape than possible with the pattern-forming interaction alone. (e-g) Inter-module dynamical independence: (e) An entire module is transiently silenced for 50 ms; (f) a large fraction of a module is externally driven by large-amplitude fixed, random, independent perturbations; (g) a continguous region that spans two modules is transiently silenced. In all cases, the perturbation remains local so neighboring regions and modules are unaffected, and the perturbed module recovers within one neural time-constant after removal of the perturbation. *See Methods for parameter and simulation details*.

We hypothesize that the second wider-scale local interaction that leads to multi-module formation might enforce pattern homogeneity even in the absence of homogeneity in the pattern-forming weights because it imposes an additional set of constraints on local pattern formation.

We numerically simulated the standard CAN model for one grid module (i.e. no gradient in the pattern forming interaction), and added noise, both in the shared radial structure of the interaction weights across neurons and i.i.d. noise in every weight, SI Fig.20 and SI sec D 9 for details and visualizations). With the same level of added noise that largely destroys patterning in conventional attractor networks (Fig. 6a, left), inclusion of a second wider-scale local interaction subject to same added noise results in robust and homogeneous pattern formation (Fig. 6a, right). We quantified this effect, finding that within-module homogeneity despite weight heterogeneity is substantially higher with peak selection than without, Fig. 6b.

The robustness enhancement from the addition of a second local interaction for single modules also holds in the formation of multiple modules (Fig. 6c). Without a second interaction, weight inhomogeneities drive variations in pattern period, whose magnitude scales like 1/*σ* (the width of the minimum of 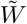) as the interaction profile is linearly stretched by *σ*, Fig. 6d (left). When a broader second scale is added, its narrower Fourier peaks confine the permitted solutions, enforcing a more homogeneous period despite weight inhomogeneity, Fig. 6d (right). Variations in period scale as ∼1/*d*, a relative reduction of *σ*/*d*. Further, since module periods scale as *d*/*m*, the smaller period modules should be more robust than larger-period modules. (Details in SI D 9).

Conceptually, the peaks of the wider local interaction “focus” the dynamics to a narrower region than specified by the narrower local interaction alone (Fig. 6d).

### Robustness to module-wide and across-module perturbations in activity

Given that the lateral neural interactions are not discontinuous across modules (weights span modules), and given that global modules emerge from local interactions and local dynamics, we wondered whether erasing patterning within entire modules or large across-module regions of the network will affect patterning in the rest of the modules. Upon silencing activity in one module (mimicking optogenetic inactivation) the other modules, module periods, and even adjacent module boundaries remain stable, Fig.6e. When a subset of cells in one module is driven to maintain randomly chosen values between 0 and 20Hz, patterned activity is disrupted in adjacent regions of the module, Fig.6f, but when the perturbation is lifted, the module period at the perturbation site is restored. If a region spanning the boundary between two modules is silenced, the dynamics and periods in spared parts of the two modules remain unchanged, and the boundary re-emerges at its pre-perturbation position after removal of the suppressive input (Fig.6g). In all cases, pre-perturbation states are restored within one cellular time-constant (≈*τ*). These findings contrast with existing models of module formation in which modules interact in a stacked architecture [61]: these models exhibit cascading dependencies between modules, so that perturbation of one module will have propagating effects across all downstream modules.

## GENERALIZED ENERGY LANDSCAPE VIEW OF MODULE EMERGENCE

Motivated by the peak-selection process of grid module emergence, we hypothesize that the same principle could supply a general mechanism for module emergence even in the absence of periodic pattern formation. The theory can be generalized in two steps. First, by translating the linear dynamics of Fourier modes into nonlinear dynamics on an general energy landscape, and next translating the Fourier peaks and troughs into multiple rugged local optima in the energy landscape (SI Sec. E).

Consider a generalized energy landscape (Lyapunov function) constructed as the sum of two terms (Fig. 7b): a spatially independent term *f* (*θ*) that is rugged with multiple similar-depth minima (constructed here as a sample from a Gaussian process with a radial basis function kernel, shown in orange in Fig. 7b) and another that has a single broad minimum (constructed here as a quadratic term, shown in green in Fig. 7b) The location of the broad minimum moves as a function *θ*_0_(*x*) of some smoothly varying parameter *x* (Fig. 7a) This results in a combined energy landscape

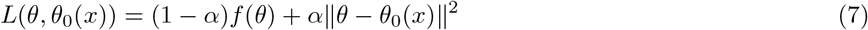

The problem can be viewed as regularized optimization on a rugged loss landscape, with a spatially-dependent regularizer ||*θ* − *θ*_0_(*x*)||^2^ acting as a prior that selects one of the local minima at each location. The dynamics of *θ* are defined as downhill flow on the Lyapunov function landscape: 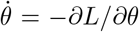. The landscape is first governed by the broad quadratic term, then sculpted by the rugged landscape (with *α* gradually decreasing with time starting from *α* = 1, SI Sec. F). Smoothly varying the parameter *x* results in a set of modular solutions with steps along the smoothly varying parameter dimension (Fig. 7c), corresponding to the state settling into the local minimum closest to the quadratic minimum at that *x*. This analytical formulation with numerical validation provides a simple mathematical framework of peak selection that shows how smooth gradients can lead to discrete patterning and modular structure emergence [31, 40, 62] beyond linear instability, Fourier modes, and periodic solutions.

**FIG. 7.**
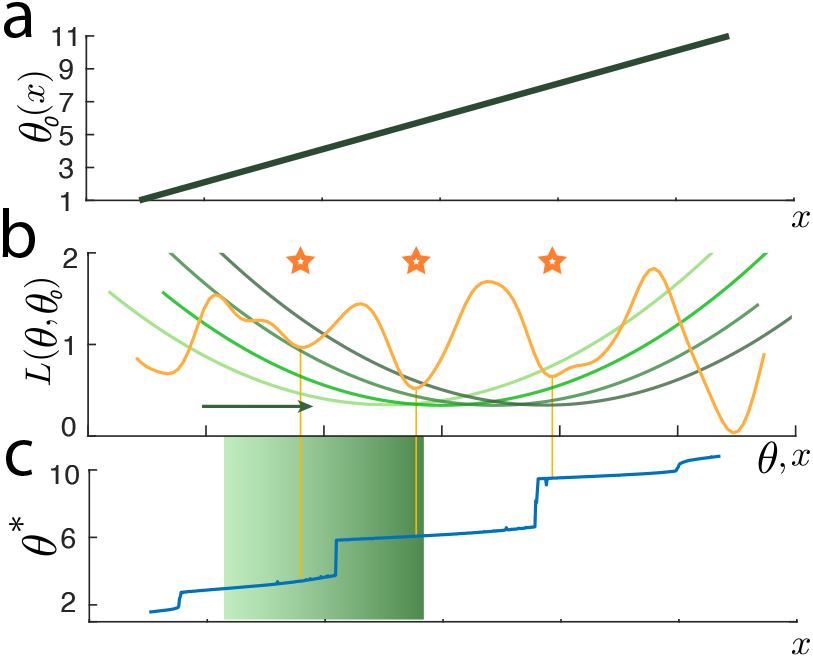
Generalized peak-selection mechanism for module emergence. (a) Parametric gradient of parameter *θ*_0_ as a function of position *x*. (b) The Lyapunov (generalized energy) function is a sum of two functions: one static function with multiple local minima, and one with a single broad minimum whose location smoothly varies with *x* according to *θ*_0_(*x*). (c) Fixed points of the dynamics upon simulation of dynamics on this Lyapunov function, as a function of the location *x. See SI Sec. F for simulation details*.

## SELF-ORGANIZATION OF ECOLOGICAL NICHES THROUGH PEAK SELECTION

The general principle of topological peak selection — with a smoothly spatially varying “selecting function” and a spatially constant fixed interaction that generates multiple local minima — can be applied to generate modularity across diverse settings. To illustrate this generality, we explore a biological example that is far from structure emergence in the brain, at a much larger scale of organization: the emergence of spatial ecological niches in ecosystems (Fig. 8a,b).

**FIG. 8.**
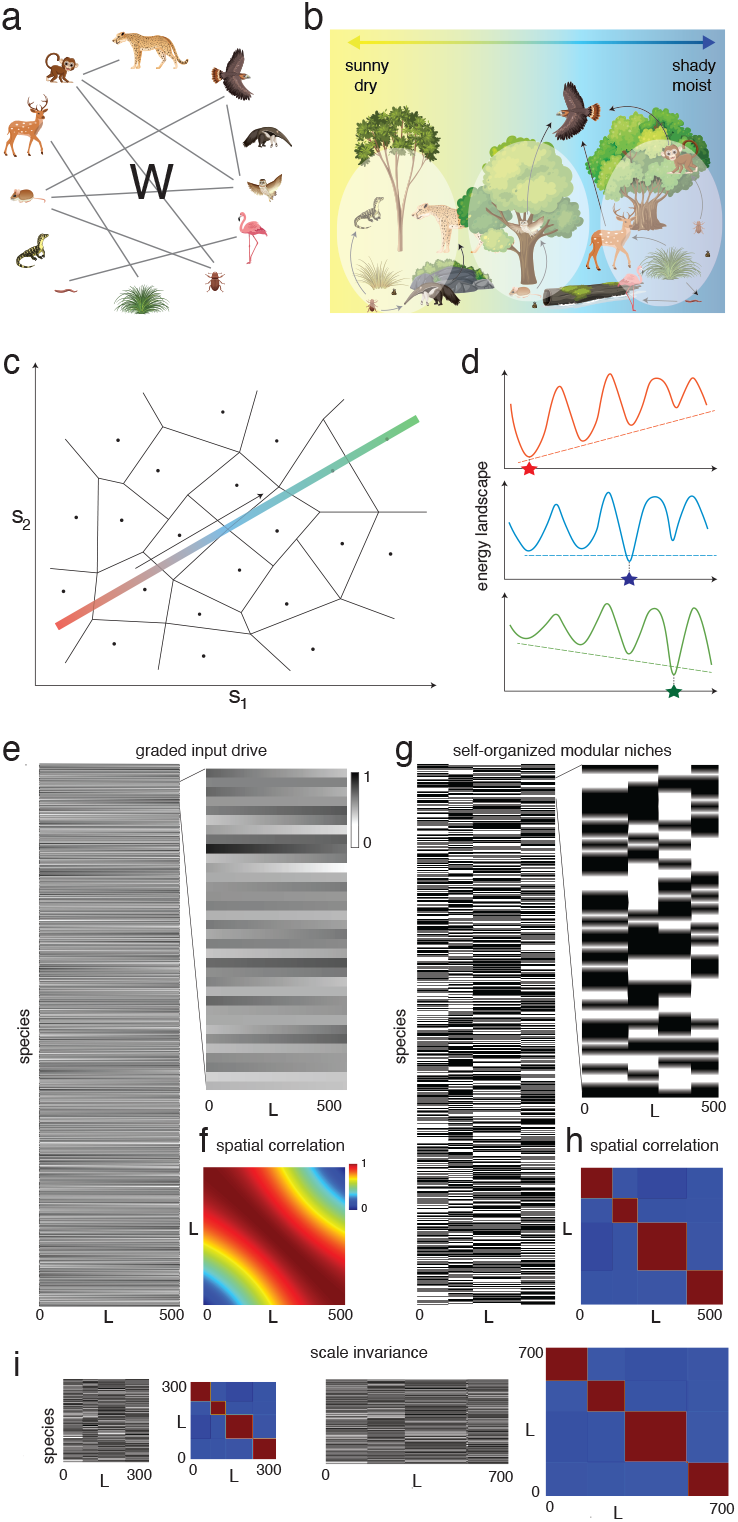
Self-organization of ecological niches through peak selection: (a) Schematic web of competitive and cooperative interactions between species. (b) Schematic of ecological niches and smoothly varying spatial resource gradients. (c) Schematic of the state space of the interacting species system as in (a), ignoring spatial distributions of species and resources, has multiple attractor states (dots; basins boundaries depicted by black lines). Multi-colored line: One-dimensional variation in resources, passing through the basins of several attractors. (d) Energy landscape of the state space visualized at 3 distinct points along the resource gradient. The resource gradient slowly “tilts” the energy landscape, gradually varying the relative heights of the local optima and discretely changing the global optimum. (e-h) Typical dynamics of the system: no spatial structure at initialization, visualized in system state (e) and spatial correlation matrix (f), self-organizes into four ecological niches at steady state (g-h). (i)Scale invariance: changing the spatial size of the system while maintaining the end points of the resource gradient function results in the same niche structure. *See Methods for parameter and simulation details*.

**FIG. 9.**
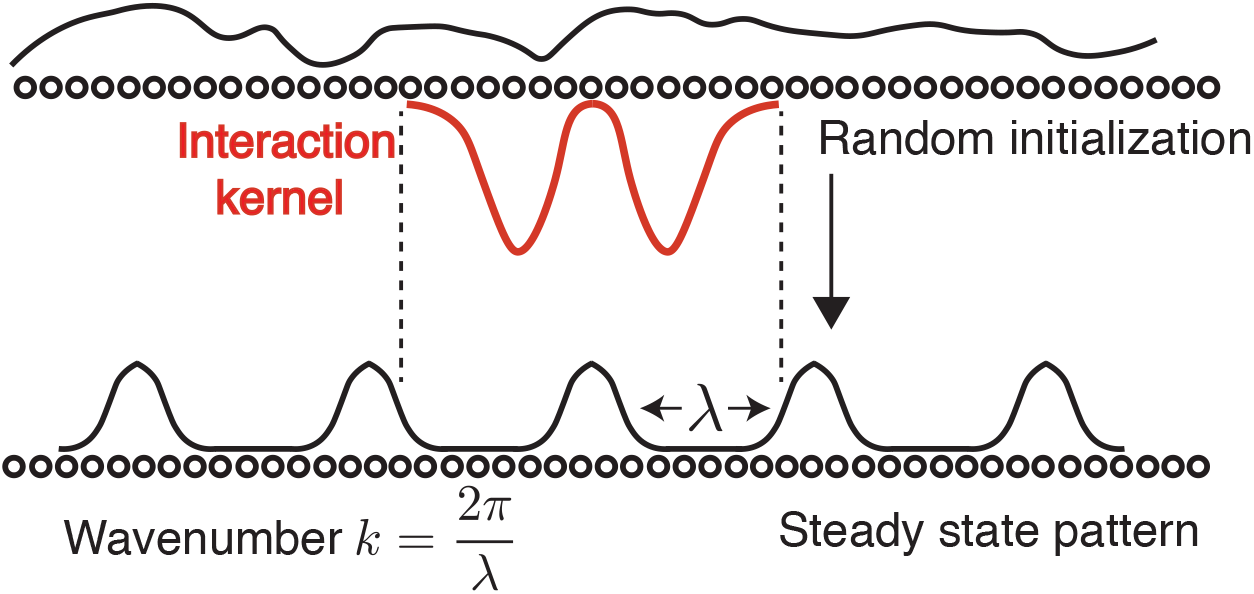
Local pattern formation in continuous attractor models of grid cells: Through local amplification of random fluctuations, the lateral interaction forms periodic patterns.

Consider a set of species that, if co-localized, interact through cooperation and competition (Fig. 8a). Each species requires different resource distributions, and resources vary along environmental gradients. Graded environmental conditions are common features of ecosystems that have been implicated in niche formation, and include for instance sunlight, temperature, precipitation, and electrochemical gradients [63–68] as well as secondary quantities like vegetation that results from these. In aquatic ecosystems, light and nutrient availability limit the quantity of primary producers, which poses fundamental limits on the food web[69, 70]. Changes in such environmental variables and nutrient availability is known to alter population dynamics[71–74] as well as inter-species interactions[68, 75, 76], that can affect the composition of the formed niche.

We model this system with Hopfield-like dynamics [77–81], where species interact according to an interaction matrix (*W*). The interaction defines a set of attractor states in a rugged energy landscape, Fig. 8c, with the minima corresponding to the set of possible niches. Interactions are also modulated by spatial distances: two species interact via *W* only if they spatially overlap. Further, species also grow in proportion to the match between the resource inputs and their own resource feature vectors. Formally, consider *N* species indexed by *i* and defined by *M* -dimensional resource feature vectors **b**_*i*_ that specify their resource needs. Each element of each resource feature vector is sampled from independent, identically distributed normal distributions (see Methods for details). A time-invariant but spatially varying resource gradient **r**^*g*^(*x*) is assumed to exist for *xϵ* [0, *L*], that is constructed by linearly interpolating between two random vectors drawn from 1, {− 1}^*M*^. The population dynamics of species *i* at position *x* and time *t* + 1 is given by:

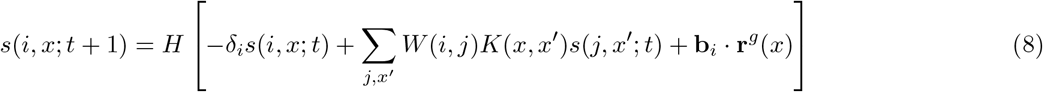

where *H*[.] is a nonlinearity, *δ*_*i*_ is the death rate, *W* (*i, j*) is a symmetric interaction matrix quantifying the cooperation and competition between species *i* and *j* (see Methods for details of construction), and *K*(*x, x*′) *≡ K*(*x* −*x*′) is a spatial interaction kernel that vanishes on a length scale much smaller than *L*. Note that while we only examine the symmetric *W* (*i, j*) case here, we expect the self-organization of modular niches to hold in general for non-symmetric *W* (*i, j*) as well. In such a case, the selection may be between different attractor states that need not be fixed point attractors.

This system settles into a steady state **s**\*. In the energy landscape view, the graded resource inputs **r**^*g*^(*x*) selectively deepen a subset of the minima. As the resource inputs continuously vary in space, the minima that are amplified vary with *x*, leading to a discrete selection of the global minima at each spatial location, Fig. 8d. We examine spatial structure through the normalized correlation matrix 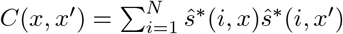, where *ŝ* is *s* normalized to have zero mean and unit norm. Continuously graded input drive with no initial modularity in the population state correlation matrix self-organizes into a steady state with modular niches (Fig.8e-h). The system also exhibits scale invariance if the resource gradient endpoints remain fixed as the spatial domain is scaled, even though the spatial interaction scale (set by *K*(*x, x′*)) is not (Fig. 8i).

## DISCUSSION

### Summary

We have combined positional and spontaneous pattern forming (or more generally, symmetry breaking) mechanisms to show that structure can robustly emerge at multiple scales from purely local interactions, without requiring modular genetic specification or modular connectivity. The resulting mechanism exhibits useful features of both processes [12]: structure is noise-tolerant and parameter invariant, self-organizing with relatively little detailed genetic instruction, and self-scaling. The mechanism involves only smooth (low information) genetically specified gradients combined with local competitive interactions whose parameters are modulated by the gradient(s), from which sharp boundaries and modules emerge.

In the context of grid cells, this is the first work to provide a framework for connecting gene expression (smoothly varying gradient(s)), structure (local neural connectivity profiles), and physiology with each other. It therefore allows future physiology, connectomics, and transcriptomics experiments to inform and constrain each other. The mechanism of module emergence generates a detailed prediction of period ratios between modules and provides a novel result on increased robustness of single continuous attractor network models with the addition of a wider local interaction.

### Predictions

The theoretical relationships – connecting simple properties of the local interactions and their spatial gradients to formed module properties – provide low-dimensional knobs for experimental testing and manipulation. The analytical and numerical models can be directly queried for predictions about the effects of varying any of the knobs. Here, we enumerate a few example predictions for connectomics (C), transcriptomics (T), development (D), and physiology (P): 1) Discrete modules are initially underpinned by merely smooth gradients in gene expression and in the interaction weights. (D,T,C) 2) The existence of an interaction that is fixed across the DV axis of MEC, and another that smoothly varies in scale (strength or width) across it. These two types of interactions may be in different cells and synapses, or the same synapses, for instance a single scaled-width weight profile with an axonal or dendritic cutoff radius that is invariant across the DV axis. (C) 3) The detailed and precise relationship of adjacent grid periods, as given by successive integer ratios, are distinct from other theoretically interestingpredictions [54, 82, 83], and deviations from successive integer ratios should be predictable by alterations in the coarse form of the (fixed-scale) lateral interaction profile (P). 4) Invariance of module number and periods to detailed local interaction shape and to the shape of the global gradient, and the shifting of module boundaries with gradient shape (T,C,P). 5)Self-scaling with system size (invariance of module number and periods despite MEC size variations if extremal values of the gradient fixed), dependence of these quantities on extremal values of the gradient, and dependence of module boundary positions on the functional form of the gradation in lateral interactions. (T,C,P) 6) Bidirectional computable relationship between lateral interaction shape and period ratios, through the scalar variable *ϕ*. (C,P) (3-6) Could be tested across individuals and species. 7) Following MEC-wide silencing, activity patterning in all modules should re-emerge independently, rather than sequentially [61]. (P) 8) Quantifiably higher robustness of dynamics to connectivity inhomogeneity, beyond the capabilities of conventional attractor models [45]. (C) 9) Independence of dynamics between modules and high robustness of dynamics to activity perturbation within and across modules: Effects of perturbation to activity are localized to the module it is applied to, without a cascading effect across modules [61]. Entirely suppressing one module should not alter others, and suppressing half a module boundary should not shift the rest of the boundary. (P) 10) General peak selection mechanism through manipulation of genetic gradients: quantitative predictions for altering the number of formed modules, periods, and positions of module boundaries with gene expression modulation. (T,D)

### Related work

This work directly extends and robustifies continuous attractor models of grid cells [45], from single modules to multiple modules, from dependence on specific interaction profiles to an infinite set of kernels for grid emergence, and from high dependence on homogeneous weights to a weaker dependence on weight homogeneity. It connects to observed DV gradients in MEC [32, 38, 42, 84–87], and also potentially more generally to observed gradients that underlie discontinuous function in cortex [31, 43]. Its focus on the theory and mechanisms of emergence of structure and function from given weights is complementary to work that models the *learning* of weights in MEC, through biologically plausible Hebbian-like rules [88] or backpropagation-based learning [83, 89–91]. In fact, the learning models generally do not produce multiple modules, and in the rare cases where they may seem to, the circuit mechanisms that produce them are unknown [89]. We know of only other work that proposes a network mechanism for multi-grid module emergence [61]. It has a distinct (stacked) initial and final architecture, and its predictions on several of the quantities noted above are interestingly and distinguishably different.

The general peak-selection principle for module emergence is both an instance and a generalization of the idea of spatial bifurcation for the emergence of discrete function from smooth gradients [31, 92]. It permits a number distinct modules to form from smooth variations in the spatial dimension, but the broader theoretical framework generalizes to variations along abstract parametric dimensions. We have shown three different flavors of peak selection-based modularity emergence: peak selection in a pattern-forming process interacting with smooth gradients in interaction parameters for grid cells; peak selection via a smoothly varying regularization term in dynamics on a rough landscape in the general Lyapunov function approach; and peak selection in a symmetry-breaking process (that is more general than Turing-like pattern formation) based on initial condition or resource gradients that tilt the landscape in a generalized Hopfield model for niche formation.

These concepts provide dynamical and mechanistic principles for *how* modular structure can emerge, in contrast to normative models that focus on *why* or *when* modular structure is favored [4, 9, 82, 93–100].

A mechanistic understanding of how discrete structure could emerge from graded precursors connects with extensive literature on the existence of smooth biophysical gradients in the brain and body [28, 31–35, 37–44, 92] even when function varies discretely.

Many works explore how structure emergence can occur with precision in the presence of noise [28, 29, 101, 102]. Some solutions within the positional hypothesis involve spatial or temporal integration of noisy gradients [12, 26, 101]. Pattern forming mechanisms have some built-in robustness to noise because the patterned state is much lower-dimensional than the overall state space [103]. Our observation that peak selection confers substantial additional robustness raises the question of whether such a mechanism might assist in tandem with positional mechanisms during developmental morphogenesis [104].

## Acknowledgements

We are grateful to Mehran Kardar for helpful discussions and to John Widloski, Akhilan Boopathy and Ling Dong for comments on the manuscript. This work was supported by the MathWorks Science Fellowship to MK, the Simons Foundation through the Simons Collaboration on the Global Brain, the ONR, and the Howard Hughes Medical Institute through the Faculty Scholars Program to IRF.

Conceptualization, writing: MK, SC, IRF; Coding, analysis: MK, SC.

## MATERIALS AND METHODS

We use a continuous attractor network (CAN) model [105–107] for grid cells [17, 18, 108, 109], with neural dynamics obeying

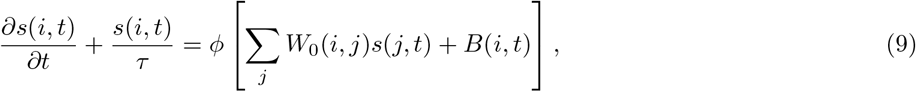

where *s*(*i, t*) represents the synaptic activation of neuron *i* at time *t, W*_0_(*i, j*) represents the synaptic strength of the coupling from neuron *j* to neuron *i, B*(*i, t*) represents the feed-forward bias to neuron *i*, and *ϕ* is a non-decreasing nonlinearity, for which we use the rectification function (*ϕ*(*z*) = [*z*]_+_ = *z* for *z* > 0 and 0 otherwise). Each neuron *i* has a preferred direction *θ*_*i*_ that is used to perform velocity integration. In the one-dimensional version of our setup, each spatial location **x** on the neural sheet has two neurons, with preferred directions *θ* = 0 and *θ* = *π*. Correspondingly, in the two-dimensional version of our setup, each location on the neural sheet has four neurons, with preferred directions *θ* = *nπ*/4 for *n ϵ* {0, 1, 2, 3}. The synaptic weights *W*_0_(*i, j*) are defined via an interaction kernel *W* (Δ*x*) such that

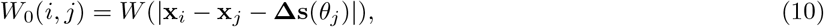

where **x**_*i*_ represents the spatial location of neuron *i*, and **l**(*θ*) is a vector with length Δ*s* oriented parallel to the angle *θ*. The feed-forward bias *B*(*i, t*; *θ*) is given by

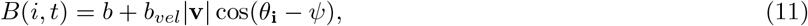

where *ψ* is the direction of the input velocity signal and |**v**| is the speed. This results in neurons with direction preference *θ* driving activity in the network towards the direction of their outgoing weight shifts **Δs**(*θ*). This mechanism is responsible for velocity integration by the network [18].

We first described the dynamics under fixed arbitrary kernels, demonstrating that they result in hexagonal pattern formation. These arbitrary kernels were constructed by interpolating between random points via the following protocol: First, we construct ‘x-values’ by considering *n* + *n*_*zero*_ uniformly spaced points from *L* to *L*, which are then perturbed by the addition of a randomly sampled number from − *L*/4*n* to − *L*/4*n* (this perturbation makes the points less regular, while disallowing consecutive points to be extremely close to each other). Second, we construct *n* ‘y-values’ sampled from a uniform distribution from −1 to 1, and define the remaining *n*_*zero*_ y-values to be 0 (the *n*_*zero*_ values at zero ensure that the interpolated function decays to zero). Then, a cubic spline interpolation (top row of Fig. 10a) or a linear interpolation (bottom row of Fig. 10a) is performed between the y-values and the x-values to generate an arbitrary function *ω*(*x*). This generated function is however not symmetric, as is required for kernel functions — thus, we construct the interaction kernel as *W* (Δ*x*) = *ω*(Δ*x*)+ *ω*(Δ*x*). Kernels whose dynamics lead to infinitely diverging firing rates are rejected and resampled. These kernels were simulated on sheets with 256 *×*256 neurons with aperiodic boundary conditions[18]. *n* was randomly chosen between 2, 3 or 4, and *L* was scaled as necessary to obtain a large number of activity bumps on the sheet to prevent finite-size effects from distorting the hexagonal lattice of activity.

**FIG. 10.**
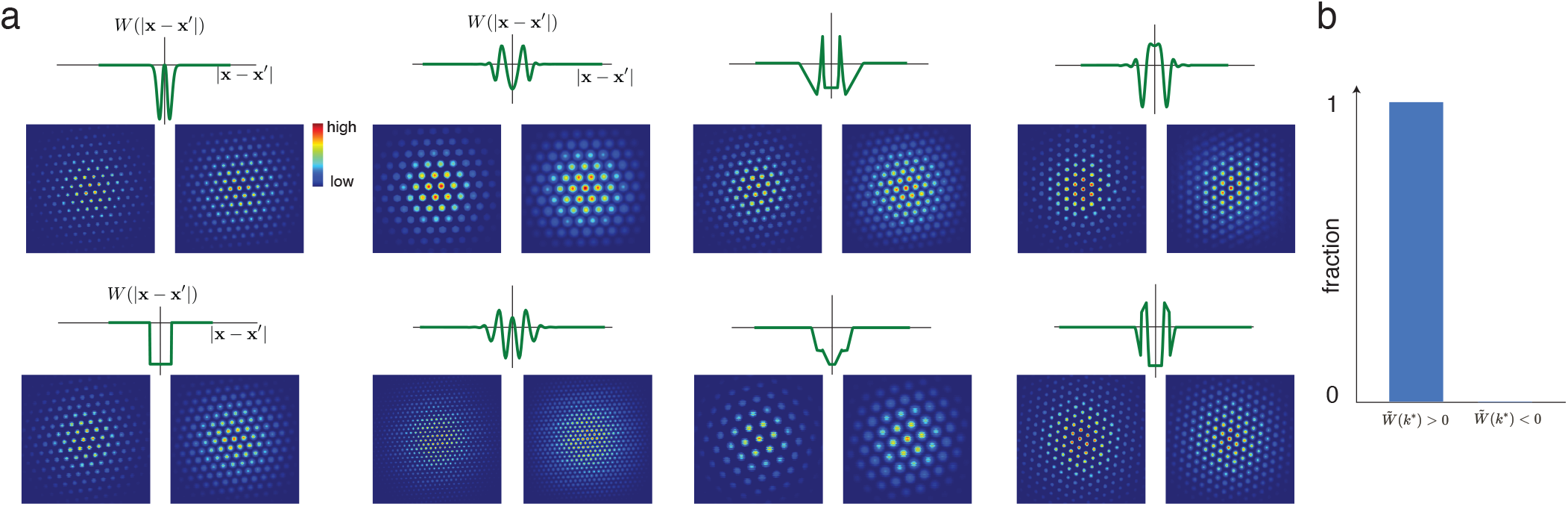
All kernels satisfying the conditions laid out in the main text can result in pattern formation, with appropriate scaling.

For the case of module formation through peak selection, the interaction weight kernel *W* is given by the sum of two components 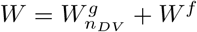 The first, 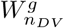 drives local pattern formation, and has a spatial scale *σ*(*n*_*DV*_), which varies smoothly in a gradient along the dorso-ventral axis, and the second, *W*^*f*^ has a fixed spatial scale *d* everywhere on the neural sheet. A variety of functions 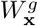 can drive local pattern formation. For concreteness, we use two specific examples: the Mexican-hat profile[18] (used in Figs. 2a-c,j,k, 11 and SI Fig. 12)

**FIG. 11.**
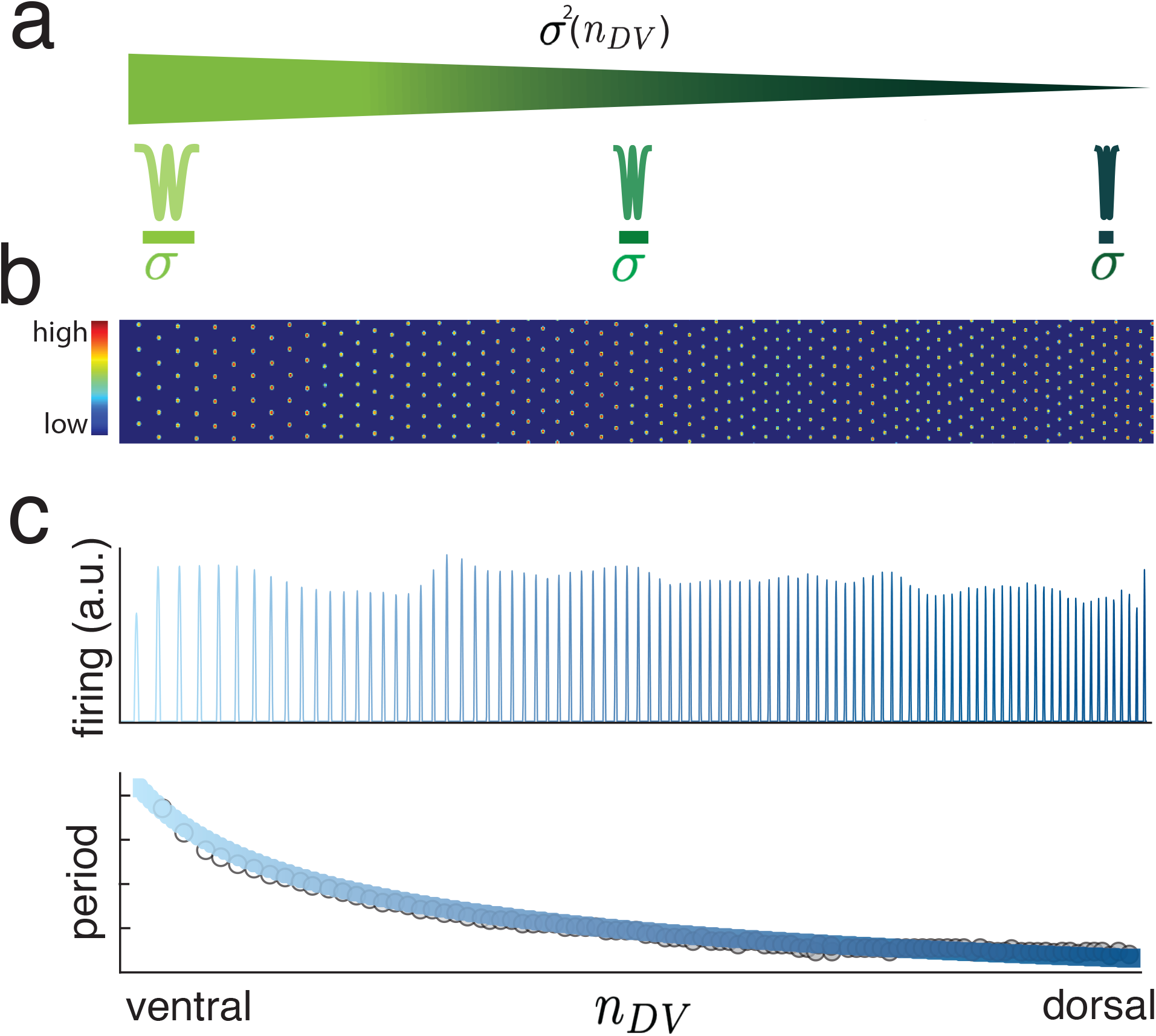
(a-c) Naive merger of the two mechanisms by smoothly scaling the width of the pattern-forming lateral interaction (j)in the grid cell CAN model [45] does not generate global modularity in 2-dimensional (b) or 1-dimensional (c) grid models: the result is one smoothly varying periodic pattern.

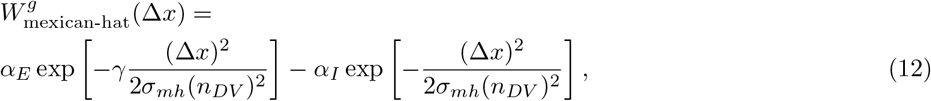

and the box-function profile[110] (used in Fig. 2 and SI Fig. 12)

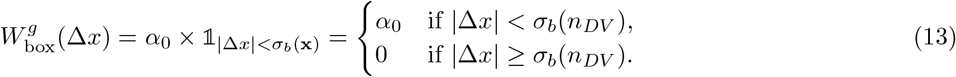

For the fixed-width interaction *W*^*f*^ (Δ*x*), we implement 3 main types — localized (used in Figs. 2,3 and SI Fig. 12), diffuse (used in Fig. 2 and SI Fig. 12) and decaying (used in SI Fig. 12).

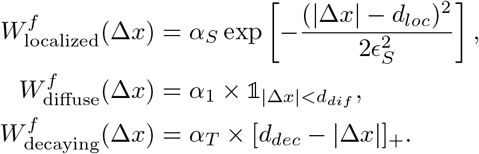

In particular,

- In Figs. 11 we use only a smoothly varying Mexican-hat pattern forming kernel 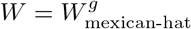
- In Figs. 2a-c,g, j,k we use 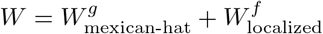
- In Figs. 2d-f,h we use a ‘Lincoln hat’ profile 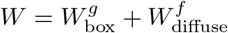
- and, in SI Fig.12 we present numerical simulations of other combinations of pattern forming and fixed-scale kernels.

To construct spatially heterogeneous kernels for analyzing the robustness to inhomogeneity in Fig. 6 we use the box function to construct

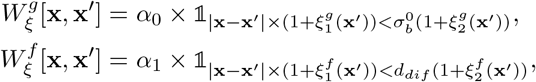

where 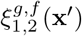 are independent random numbers chosen uniformly from 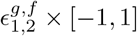 For the particulars of Fig. 6b, 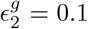 and other noise terms are set to zero (In the one-dimensional case *ϵ*_1_ and *ϵ*_2_ have the same effect); for Fig. 6c, 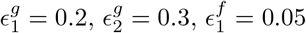 and 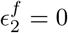 (See SI Sec. D9 for more details).

Table I we present a list of common parameters used across all numerical grid-cell simulations. Then, in Tables In II,III we present the parameter values used for the kernels used in our numerical simulations

**TABLE I.** Parameters held constant across all numerical simulations.

| Parameter | Value |
| --- | --- |
| $\tau$ | 30 |
| $dt$ | 0.05 |
| $b$ | $\begin{cases} 70 & \text{in 1D} \\ 1 & \text{in 2D} \end{cases}$ |
| $b_{vel}$ | $\begin{cases} 105 & \text{in 1D} \\ 1 & \text{in 2D} \end{cases}$ |
| $\Delta s$ | 2 |

**TABLE II.** Pattern forming kernel parameters used for numerical simulations.

| $W_{\text{mexican-hat}}^g$ parameters | Value |
| --- | --- |
| $\alpha_E$ | 1000 |
| $\alpha_I$ | 1000 |
| $\gamma$ | 1.05 |
| $N^{1D}$ | 3000 |
| $N_y^{2D}$ | 100 |
| $N_x^{2D}$ | 1000 |
| $\sigma_{mh}(n_{DV})$ | $1/\sqrt{2\beta(n_{DV})}$ |
| $\beta(n_{DV})$ | $\beta_0 + (\beta_1 - \beta_0)n_{DV}/N'$ |
| $N'$ | $\begin{cases} N^{1D} & \text{in 1D} \\ N_x^{2D} & \text{in 2D} \end{cases}$ |
| $\beta_0$ | $\begin{cases} 2.5 \times 10^{-2} & \text{in 1D} \\ 3/676 & \text{in 2D} \end{cases}$ |
| $\beta_1$ | $\begin{cases} 2.5 \times 10^{-1} & \text{in 1D} \\ 9/338 & \text{in 2D} \end{cases}$ |
| $W_{\text{box}}^g$ parameters | Value |
| $N^{1D}$ | 5000 |
| $\alpha_0$ | -40 |
| $\sigma_b(n_{DV})$ | $15 + 30n_{DV}/N$ |

**TABLE III.** Fixed-scale kernel parameters used for numerical simulations.

| $W_{\text{localized}}^f$ parameters | Value |
| --- | --- |
| $\alpha_S$ | 4 |
| $d_{loc}$ | $\begin{cases} 84 & \text{in 1D} \\ 50 & \text{in 2D} \end{cases}$ |
| $\epsilon_S$ | $\begin{cases} 4.77 & \text{in 1D} \\ 1.6 & \text{in 2D} \end{cases}$ |
| $W_{\text{diffuse}}^f$ parameters | Value |
| $\alpha_{dif}$ | -0.25 |
| $d_{dif}$ | 135 |
| $W_{\text{decaying}}^f$ parameters | Value |
| $\alpha_T$ | 25 |
| $d_{dec}$ | 150 |

For the modular niche formation, we consider the setup as described in Eq. 8, with *N* = 1000 species, each characterized by a random *M* = 2000 dimensional random feature vector indicating resource preference. We numerically simulate our setup on a discrete lattice *x* ∈ {0, *L*} for *L* = 300 in Fig. 8i left, and *L* = 500 otherwise.We instantiate the nonlinearity *H* as a shifted Heaviside function, *H*[*x*] = 1 for *x* ≥ 0.5, and *H*[*x*] = 0 otherwise, and choose the death rate *δ*_*i*_ = 0.1 for all species. To construct *W* (*i, j*) as an interaction matrix that quantifies the cooperation and competition between species, we follow a set up similar to a Hopfield model with {0, 1} activity[111]. We first choose a set of random points in *N* -dimensional species space **s**_*q*_ for *q* ∈ {1, …, *Q*}, denoting *potential* niches. We choose *Q* such that 1 ≪ *Q* ≪ *N*. Each **s**_*q*_ vectors consists of a +1 at elements corresponding to species that may co-exist, and −1 otherwise. In practice, we draw each element uniformly from the set {0, 1}, constructing an *N ×Q* matrix. The weight matrix *W* (*i, j*) is then constructed as

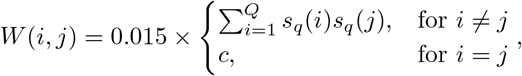

where *c* is a positive constant set to 10.

The spatial interaction kernel *K*(*x, x*′) = *K*(*x* –*x*′) is chosen to be a Gaussian function with standard deviation 1.75 (which is much smaller than the entire spatial extent of the system, *L*). The end points of the resource gradient are chosen as two random *M* = 2000 dimensional vectors with elements draw independently from i.i.d. Gaussian distributions with zero mean and standard deviation 2/*N*, and the preference vectors **b**_*i*_ are drawn from i.i.d. Gaussian distributions with zero mean and unit standard deviation.

The initial condition for the simulation is set to be the uniform state *s*(*i*) = 0.5 for all *i*, and the simulation is run until the dynamics reach a fixed point state. The final formed fixed point state is examined by calculating the correlation matrix

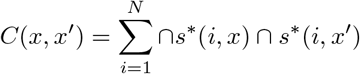

where

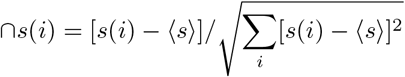

where ⟨*s*⟩ = (1/*N*) Σ_*j*_ *s*(*j*).

## SUPPLEMENTARY TEXT

The supplemental information is structured as follows: First, in SI Sec. A we present the mathematical analysis for pattern formation, and generalize the theory of CAN models of grid cells to show analytically and numerically that an infinite set of local interaction kernels can generate a grid cell network, as shown in Fig. 2 and Fig. 10. Second, we demonstrate analytically and numerically in SI Sec. B that simply introducing a gradient in the pattern forming kernel of the continuous attractor model is *not* sufficient to result in modularization, as demonstrated in Fig. 1 of the main text. Third, in Sec. B 1 we show how the addition of a Gaussian localized kernel results in self-organized modularization. Fourth, we show in Sec. D that among arbitrary kernels, those with simple shapes result in a simple equation describing the detailed period ratios of the formed grid modules as shown in Fig. 4. Fifth, this will lead to simple estimates for the number of modules and their sizes in terms of other system parameters, which we derive in SI Sec. D7. Sixth, after having described our results primarily for the case of one-dimensional grid cells, we then demonstrate in Sec. D8 that our arguments extend naturally to two dimensions, and we present numerical results demonstrating the same. Seventh, in SI Sec. D10 we then demonstrate that our results and predictions of grid period ratios are consistent with available data sources to a large extent. Finally, we generalize our result to the context of dynamics on a rough energy landscape (SI Sec. E), and provide broader perspectives of our results in the contexts of general loss optimization (Sec. F) and eigenvector localization (SI Sec. G).

## Appendix A Generalization of grid cell CAN dynamics theory: infinite set of interactions produce grid cells

It is known that Mexican hat-like kernels [18] and Lincoln hat-style kernels [48] generate grid patterning. While there are analytical results on why grid patterning emerges from a Mexican hat interaction, the Lincoln hat result is empirical, without theory. Here we seek to explain when grid patterning emerges, and to determine other kernel shapes that are consistent with it.

Consider the standard equations for the dynamics of recurrently connected neurons (expressed for notational simplicity in the continuum or large neural number limit):

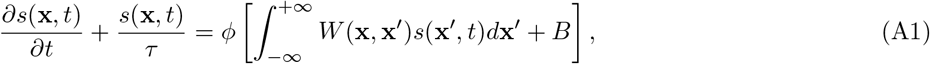

where *s*(**x**) is the synaptic activation of the neuron at the vector position **x** on a 2-dimensional neural sheet, *W* (**x, x**′) is coupling strength from a neuron at **x**′ to a neuron at **x**, *τ* is the biophysical time-constant of individual neurons, *ϕ* is a non-negative monotonic transfer function, and *B* is a uniform feedforward input to all neurons. The neural nonlinearity is anything non-symmetric (*ϕ*(*x*) = *ϕ*(−*x*), for reasons given below and in [112–114]. For simplicity, we select the rectification function (*ϕ*(*z*) = [*z*]_+_ = *z* for *z* > 0 and 0 otherwise.

From the linear (in)stability analysis, four conditions on the interaction kernel weights *W* (**x, x**′) may be sufficient for grid-like patterning: 1) For global stability, let *W* (**x, x**′)*d***x** < 0 (this is consistent with models of grid cells with negative recurrent coupling [45, 110] and with experiments suggesting that grid pattern formation might dominated by recurrent inhibitory circuitry [110]). 2) Let the interactions be radially and translationally symmetric, *W* (**x, x′**) *≡ W* (|**x** − **x** |; *σ*), which means that the Fourier transform can be written in terms of its radial part: 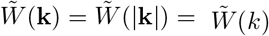. 3-4) To ensure a non-zero wavelength *k* of pattern emergence, the Fourier transform of *W* should satisfy that its maximum occurs at a non-zero value of *k, k*\* = arg max 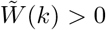, and that this maximum should be positive and sufficiently large, 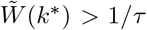. Note that conditions 3-4) can be easily made true so long as *W* is not everywhere negative, and we are permitted a global scaling factor to ensure that the positive component is sufficiently large.

The emergent activity pattern will consist of superpositions of waves with period 2*π*/*k*\* [14, 18, 112, 115–117]. This period scales as *σ*, the characteristic width of the interaction kernel *W*. The specific geometry of the emergent period-2*π*/*k*\* pattern depends on the relative strengths and interactions of the waves of wavenumber *k*\*. If the interaction kernel is isotropic and the boundary conditions are infinite or isotropic, the formed pattern will be an equally-weighted superposition of all three waves of wavenumber *k*\*, defining a triangular lattice. The phase of the formed pattern will be set by spontaneous symmetry breaking.

The non-symmetric nature of the transfer function results in patterns with *hexagonal* rather than other symmetries [112–114] because XXX The set of conditions above are hypothesized to be sufficient for grid-like patterning.

How many kernel functions *W* satisfy these conditions? Essentially, an infinite set does so (with rare exceptions). First we discuss some of the exceptions to gain some insight. Gaussian and Lorentzian functions, when they are positive, have a single peak in their Fourier transforms at *k* = 0 when the functions are positive. When the functions are negative everywhere, they fail to satisfy condition 1). Thus, Gaussian and Lorentzian functions are two special functions that do not satisfy the criteria 1)-4). However, as argued in Sec. C, making small perturbations to functions that do not satisfy 1)-4) results in the conditions 1)-4) being satisfied, suggesting that the functions that do not satisfy 1)-4) are a small and very special set, and that most functions can be scaled to satisfy 1)-4).

We next performed numerical experiments to test the hypothesis that randomly generated functions will generically have Fourier Transforms that are not negative everywhere or only non-negative at 0, and therefore might generate grid-like patterning (see Methods for details of random sampling of kernel functions). We found that indeed randomly constructed kernel functions satisfied the hypothesized property for their Fourier transforms: we generated 10^6^ random localized kernel functions, and all of these satisfied the conditions of being not negative everywhere or being non-negative only at *k* = 0 (SI Fig.10). We further found that these kernel functions, under the further condition that they did not produce diverging neural activity, generated hexagonal patterns. Some of these are shown in Fig. 2a. In sum, an infinite set of local interaction profiles will generate grid cell-like activation patterns. Such candidate profiles can be generated at random and with very high probability generate grid-like patterning.

## Appendix B Pattern formation with graded kernels

Motivated by the experimental observations described in the main text, we modify the Mexican-hat function to introduce a smooth gradient in the characteristic interaction widths *σ*_*E*_, *σ*_*I*_.

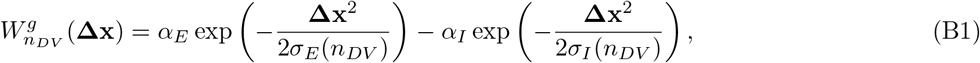

where *σ*_*E*_(*n*_*DV*_) and *σ*_*I*_ (*n*_*DV*_) are now functions that depend on position in the neural sheet, and encode the smoothly varying characteristic scale of the Mexican-hat interaction along the dorso-ventral axis:

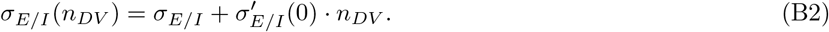

For such graded kernels, we will use *W*(x, x′) and 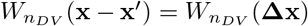 interchangeably. In this case, Eq. A1 then becomes

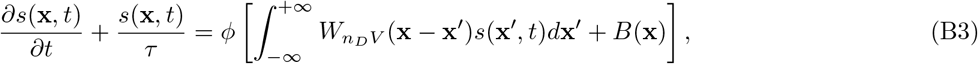

Under this approximation, we perform a linear stability analysis of the neural dynamics, to identify the the growing periodic modes locally at the position on the neural sheet *n*_*DV*_.

We first identify an unstable steady-state solution to Eq. (B3), which we denote as *s*_0_(**x**). This solution satisfies

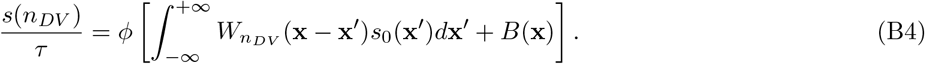

In the limit of very slowly varying changes in 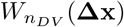 as a function of *n*_*DV*_, the unstable steady state solution will be

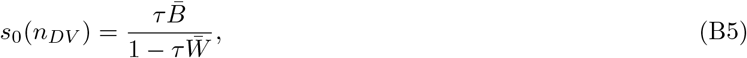

where 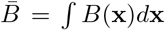 and 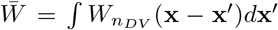, (For 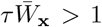 the only locally homogeneous steady state is *s*_0_(*n*_*DV*_) = 0 due to the rectifying nonlinearity, which as we justify shortly cannot support periodic pattern formation due to being a stable fixed point).

We then consider a perturbative analysis, by examining the evolution of *s*(**x**, *t*) = *s*_0_(*n*_*DV*_)+ *E*(**x**, *t*). We apply our analysis to the early time evolution of this initial condition, such that *ϵ*(**x**, *t*) ≪ *s*_0_(*n*_*DV*_). Inserting our form of *s*(**x**, *t*) in Eq. (B3), we obtain

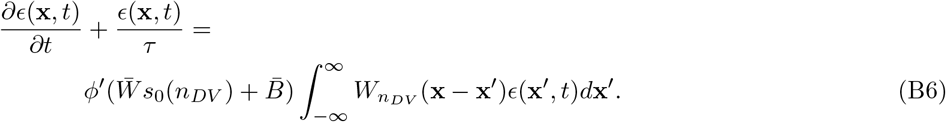

Since 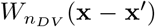 is a local kernel, we approximate the above integral with one evaluated over the region{**x′** :|**x** – **x′** |< *l*}, with *l* much larger than the length-scale of the kernel 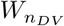 at all **x**. Over this interval, we posit that *ϵ*(**x**′, *t*) = *ϵe*^*i***k**·**x′**+*α*(**k**)*t*^, where *α*(**k**) denotes the growth rate of this *ϵ* perturbation. Inserting this form into Eq. (B6) yields,

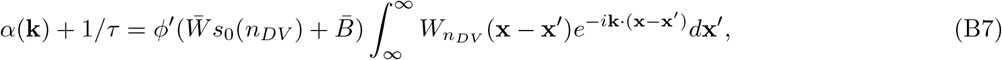

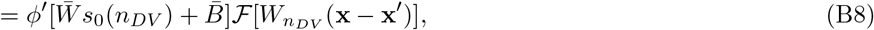

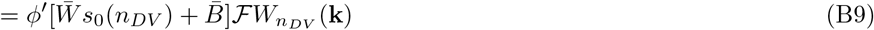

where 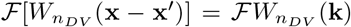 is the Fourier transform of the interaction kernel corresponding to position *n*_*DV*_ on the neural sheet. For the rectifying nonlinearity *ϕ′* = 1, and the requirement for the periodic perturbation to be growing is 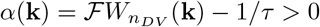

Note that since 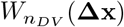 is a kernel, it is a radially-symmetric real function, and hence the Fourier transform 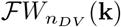 will also be real function that is radially-symmetric in *k*. Thus, for simplicity, we will only focus on the magnitude of **k**, which we denote as *k* = |**k**| ≥ 0 (In this context, for the two-dimensional case, one may re-interpret the radial component of the Fourier transform of 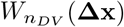 as the Hankel transform of 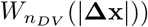

By definition, the magnitude of the wave vector *k*\* that corresponds to the fastest growing mode locally around position **x** on the neural sheet will be the **k** that maximizes *α*(**k**). Under the approximation of slow changes in the length-scale of the interaction kernel 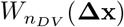, we see from Eq. (B9) that

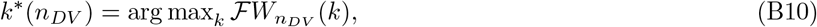

since 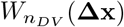 (and hence *s*_0_(*n*_*DV*_)) has been assumed to have a negligible dependence on *n*_*DV*_.

For 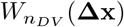 given by Eq. (B1), i.e., without any additional fixed-scale interaction, we obtain from Eq. (B10)

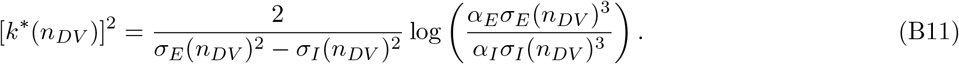

If we assume that *σ*_*E*/*I*_ (**x**) = *η*_*E*/*I*_*σ*(*n*_*DV*_), where *η*_*E*_ and *η*_*I*_ are **x**-independent constants, then we obtain

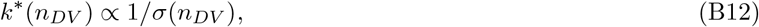

and hence

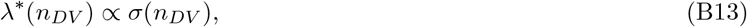

where *λ*\*(*n*_*DV*_) is the periodicity of the grid pattern formed locally around position *n*_*DV*_. This results in a smooth change of grid period, corresponding to the observation in Fig. 1g of the main text.

Note that this result is generally true for any pattern forming kernel 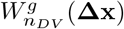 that has a Fourier transform with at least one local maximum, and *does not* rely on the specific form of a Mexican-hat interaction. Indeed, Eq. (B13) holds for any kernel 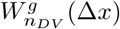 that depends on a length-scale *σ*(*n*_*DV*_). As an example, we present the corresponding analysis for the box-shaped kernel employed for pattern formation in Ref. [110].

In this case

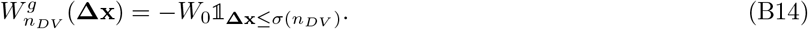

As discussed above, the quantity of interest is 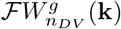

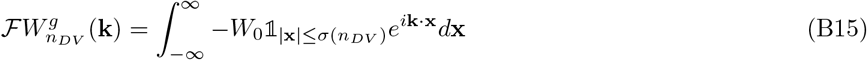

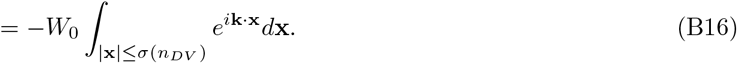

The above integral can be calculated in a one-dimensional setup to obtain

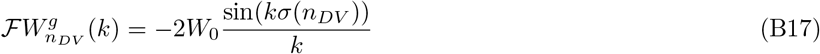

and can be calculated in a two-dimensional setup to obtain

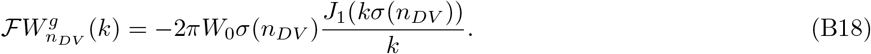

In both of the above cases, note that *k*\* ∝ 1/*σ*(*n*_*DV*_) since *σ*(*n*_*DV*_) is the only length-scale characterizing the kernel 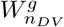. In particular, numerical maximization yields

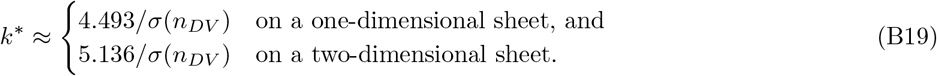

### 1. Fixed-scale interactions and modularization

We now claim that the addition of a fixed-scale kernel, *W*^*f*^ (**Δx**) is sufficient to result in modularization of grid periods, with discrete changes in grid period as a function of spatial position along the dorso-ventral axis. This set of interactions can effectively be implemented by two populations of interneurons - one with fixed arborization and weaker synaptic connections and one with varying arborization length and stronger synaptic connections.

For simplicity, we shall present the specific Fourier transform computations for the one-dimensional problem, although we note that all of the qualitative results hold in two dimensions as well, with the Fourier transforms of the relevant functions replaced with their Hankel transforms (as shown in Sec. D8).

We include an additional weak interaction term *W*^*f*^ that critically does *not* depend on the neural sheet position *x*. For reasons that will become apparent soon, we choose kernels *W*^*f*^ (Δ*x*) such that the Fourier transform changes sign a sufficiently large number of times. We hypothesize that this requirement is not particularly restrictive, and will demonstrate that this holds for most kernels *W*^*f*^.

The entire interaction profile is then given by

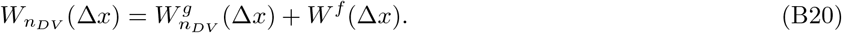

We first demonstrate our result with an example of a simple kernel, to justify how Eq. (B10) leads to the emergence of discrete grid modules. Consider the localized excitatory interaction

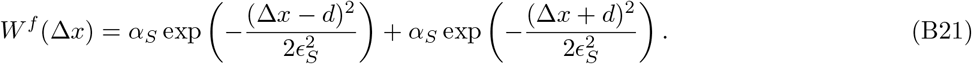

Corresponding to our interpretation of *W*^*f*^ (Δ*x*) above being a localized kernel, we choose *E*_*S*_ ≪ *d*.

This choice of 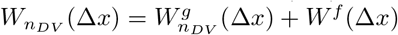 leads to the the Fourier transform,

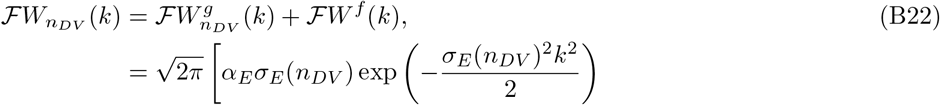

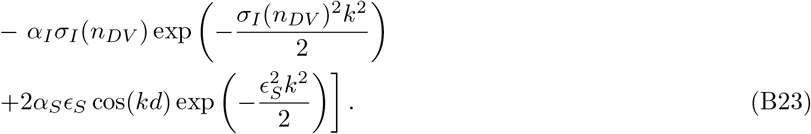

In our model, the magnitude of the *W*^*f*^ (Δ*x*), i.e., *α*_*S*_, is chosen to be smaller than the magnitude of the Mexican-hat interaction. Thus we interpret ℱ*W*^*f*^ (*k*) in Eq. (B23) as being a small perturbation to the Fourier transform of the standard Mexican-hat interaction, 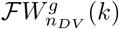 Further, since *d* is assumed to be much larger than the scale of the Mexican-hat, *σ*_*E*/*I*_, then the term cos(*kd*) in ℱ*W*^*f*^ (*k*) oscillates at a *k*-scale much smaller than the relevant scales of 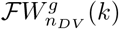 (see Fig. 3b-c of the main text). Additionally, since *E*_*S*_ ≪ *d*, the gaussian envelope multiplying the rapidly oscillating term has a scale 1/*E*, which is much larger than the periodicity 1/*d*.

Thus, in *k*-space, the rapidly oscillating term, ℱ*W*^*f*^ (*k*) can be thought of as predefining a set *S* = {*k*_1_, *k*_2_,…} of local maxima. Under the approximations made above, the addition of the smoother function 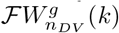 will not change the position of the local maxima. This results in the *local* maxima of 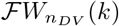 also being the same set *S*. Importantly, we note that since *S* was predefined purely via ℱ*W*^*f*^ (*k*), *there is no n*_*DV*_ *dependence on the set S*.

Following Eq. (B10), the wave-vector corresponding to the pattern formation at point *x* on the neural sheet corresponds to the *global* maxima of 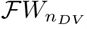 (*k*). Thus, at all points, the pattern formation corresponds to one of the discrete set of choices of wave vectors, *S* = {*k*_1_, *k*_2_ .. .}. As can be seen from Fig. 3c, the smoothly varying gradient in the Mexican-hat term, 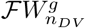 as a function of *x* picks different choices of *k*_*i*_ depending on the position *n*_*DV*_ — the *k* ∈ *S* that is nearest to the maxima of 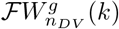 will be chosen as the global maxima, and will be the wave vector corresponding to the pattern at *n*_*DV*_. We refer to this mechanism as “peak selection”.

For our particular choice of *W*^*f*^ (*x*) made in Eq. (B21), we obtained

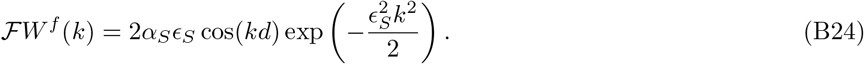

We can then approximate the local maxima of ℱ*W*^*f*^ (*k*) as occurring at

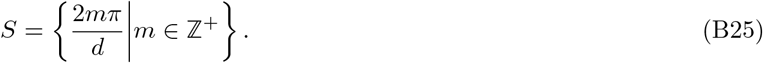

This immediately indicates that the ratios of periods of successive grid modules will be given by

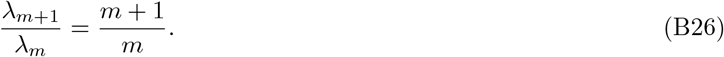

Thus, the addition of a fixed-scale interaction, *W*^*f*^ such as Eq. (B21) results in discrete grid modules. We now show that this peak-selection mechanism, and hence modularization, occurs for arbitrary choices of the fixed-scale interaction kernel *W*^*f*^ (Δ*x*).

## Appendix C Kernels that lead to modularization

The peak-selection modularization mechanism described above arises naturally from the presence of the rapidly oscillating term in ℱ*W*^*f*^ (*k*). In fact, for discrete grid modules to occur, the only constraints imposed on the fixed-scale kernel *W*^*f*^ are: (a) the Fourier transform ℱ*W*^*f*^ (*k*) must have a sufficiently large number of maxima (at least 4 maxima, corresponding to the 4 grid modules observed in experimental observations); and, (b) these maxima must be at scales smaller than 1/*σ* in *k*-space. Here we argue that this is generally true for arbitrary kernels, modulo a single scaling parameter.

We hypothesize and give support, without formal proof, that almost every arbitrarily chosen kernel *W*^*f*^ (Δ*x*) will have a Fourier transform with multiple maxima satisfying condition (a). We will then argue that this kernel can always be scaled to satisfy condition (b).

To motivate our hypothesis, we first note that it is actually possible to construct specific kernels *W*^*f*^ (Δ*x*) whose Fourier transform does not present multiple maxima. For example, the Gaussian kernel, *W*_*gauss*_(Δ*x*) = exp[(−Δ*x*)^2^/2], results in a Fourier transform that is unimodal. However, we hypothesize that such functions are rare in the space of all continuous functions in *L*^2^. Indeed, we can construct a function that is arbitrarily close to the Gaussian kernel whose Fourier transform will have an infinite number of maxima: Let *f*_0_(Δ*x*) = 1_[−1,1]_ be the box function. Define

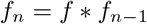

for all *n* ≥ 1, where *f* * *g* represents the convolution of functions *f* and *g*. By the central limit theorem,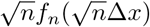 will approach *W*_*gauss*_(Δ*x*). However,

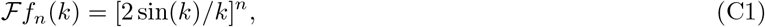

which clearly has an infinite number of maxima. Thus,even though the Gaussian kernel has a unimodal Fourier transform, we can construct a function 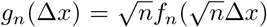 that is arbitrarily close to the Gaussian kernel (for sufficiently large *n*) but has a Fourier transform that presents an infinite number of maxima.

In this context, we claim that almost every arbitrarily chosen kernel *W*^*f*^ (Δ*x*) will have a Fourier transform with multiple maxima. This may be intuited as follows: First note that Fourier space is a dual space, and hence instead of considering arbitrary kernels in real space we may equivalently choose arbitrary kernels in Fourier space. Further assuming that ℱ*W*^*f*^ (*k*) is a smooth function, we hypothesize that generically smooth functions that are in *L*^2^ will almost always have multiple maxima and minima. Note that this heuristic also applies to the pattern forming kernel as well — we hypothesize that generic *L*^2^ smooth functions will have some maxima and minima with a global maxima that exists at *k* > 0 with probability 1, and will not be always negative (in which case a rescaling will make the maxima larger than the constant specified by requirement 2 for pattern forming kernels in the main text). Thus we expect that kernels will generically result in hexagonal pattern formation, as demonstrated in Fig. 10.

Thus condition (a) may be satisfied for arbitrary kernels *W*^*f*^ (Δ*x*).

Next, note that scaling a function in real space results in an inverse scaling of the Fourier transform, i.e., ℱ[*W*^*f*^ (*a*Δ*x*) = ℱ*W*^*f*^ (*k*/*a*). Hence, we can always scale the function *W*^*f*^ (Δ*x*) to obtain a Fourier transform with maxima that are within any desired scale, allowing condition (b) to be satisfied.

In Fig. 12, we show examples of modularization arising from different combinations of graded pattern forming kernels (*W*^*g*^) and fixed-scale kernels (*W*^*f*^). In each case, we also present the expected periodicity in each module as a function of spatial position as given by the perturbative analysis Eq. (B10). The analytical result based on linear stability provides an excellent prediction of the pattern periods per module (see also Main text, Fig. 3e). It also predicts the locations of the module boundaries (see also Main text, Fig. 3e) though not as accurately: module boundary predictions tend to be slightly but systematically offset relative to the simulated dynamics, due to the effects of nonlinearity in the later stages of pattern formation.

**FIG. 12.**
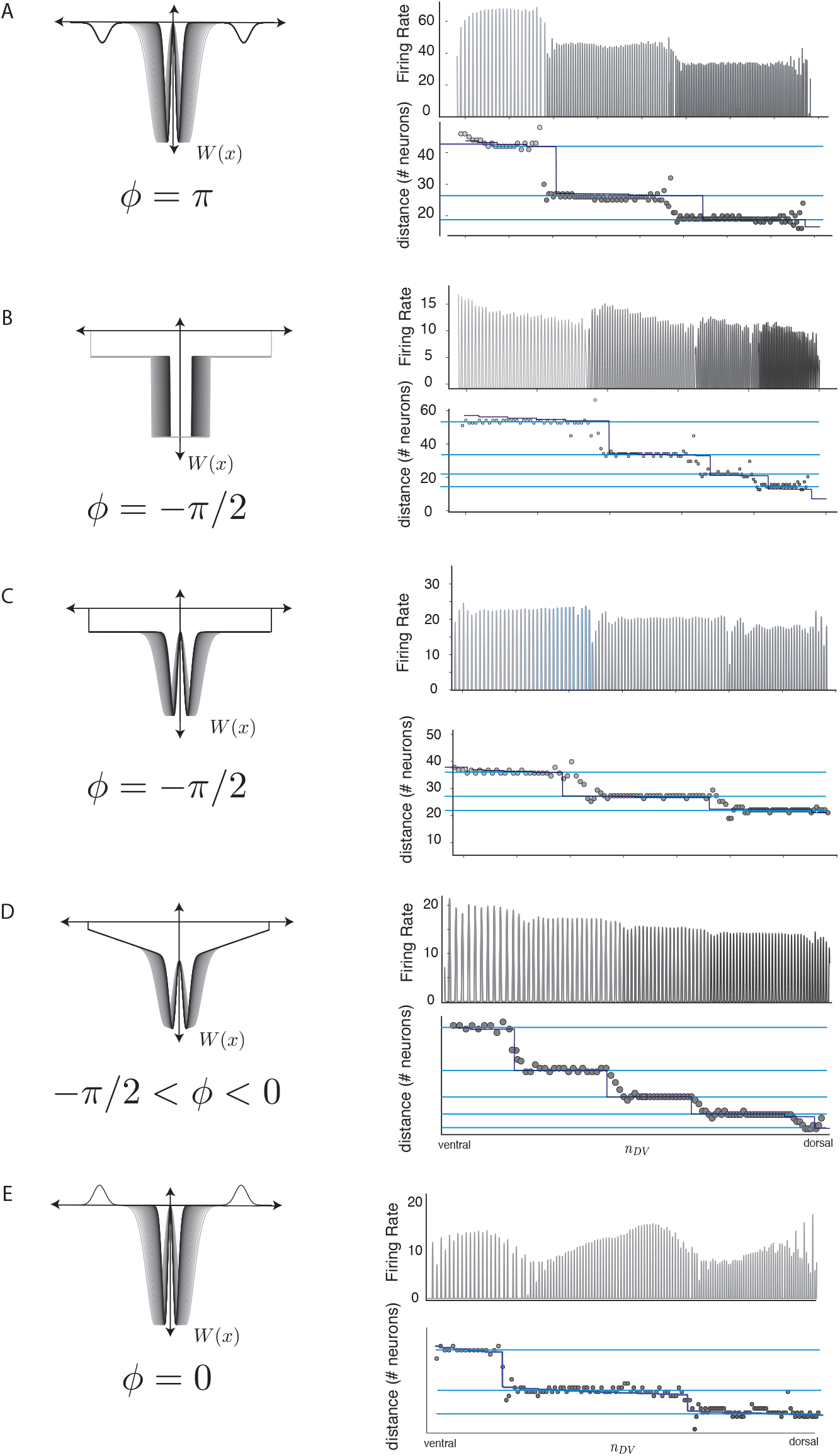
Examples of modularization and population activity (right column) with various pattern forming and fixed-scale lateral interactions (left column). In each case the dark-blue curve shows the predicted value of the grid period from Eq. (B10), and is in close agreement with the numerical simulation of the population activity. Each of the fixed-scale interactions has a qualitatively different shape, spanning different values of *ϕ* (see Fig. 3)

**FIG. 13.**
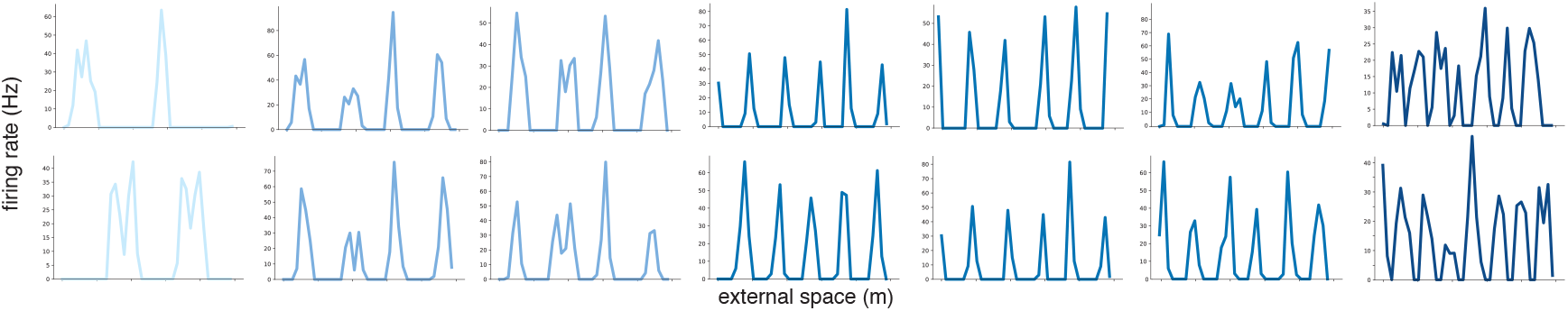
Sample tuning curves from several neurons in all modules from the network of Fig 2a.

**FIG. 14.**
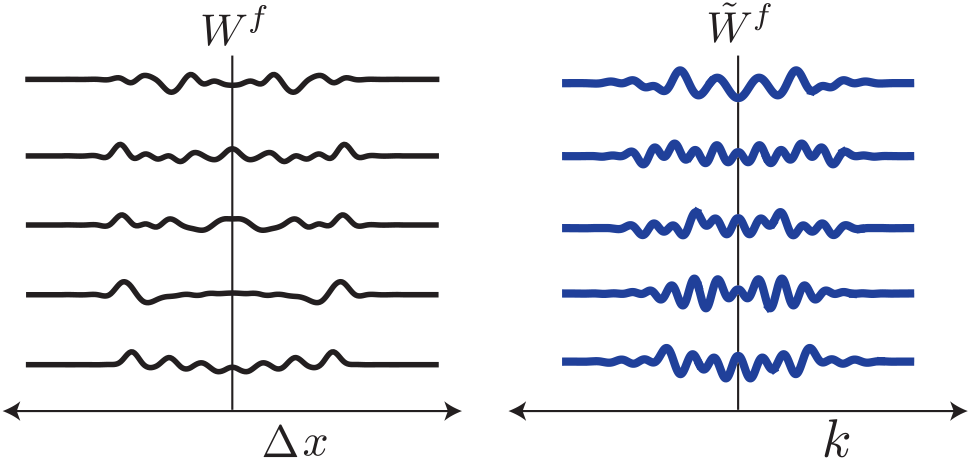
Randomly constructed fixed-scale interactions (left column) and their Fourier transforms (right column), in addition to the hand-designed ones in Fig.3, that give *ϕ* = 0 .

## Appendix D Simple kernels and period ratios

What kinds of fixed-scale interactions might be present in the medial-entorhinal cortex? As described in the main text, in the context of biology, we might expect *simple* interaction kernels *W*^*f*^ to be relevant i.e., the fixed-scale interaction profile *W*^*f*^ has the following characteristics: (a) there exists a *single* length-scale *d* that primarily characterizes the shape of *W*^*f*^ ; (b) any other length-scales relevant to *W*^*f*^, say scales *ϵ*_1_, *ϵ*_2_, … are each much smaller than the primary length scale *d*. Further, we assume that the primary length-scale associated with the fixed-scale interaction is larger than the length-scales of the pattern forming kernel, i.e., *d* ≫ *σ*_*E*/*I*_ (*n*_*DV*_).

We will demonstrate that *simple* fixed-scaled interaction kernels result in analytic expressions for grid periods that are characterized by a single angular variable *ϕ*

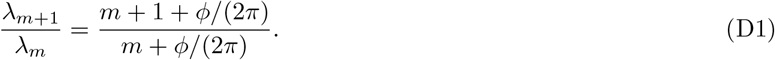

Before filling in the details of our argument, we present an intuitive explanation of the general idea:

Consider the following basic classes of *simple* kernels that satisfy the above-described criteria corresponding to a length-scale *d*:

1. *g*(|Δ*x*| − *d*), for arbitrary functions *g*(*ρ*) that are nonzero only over scales |*ρ*| < *ϵ*_*i*_ (a *localized* kernel), and,
2. A constant term, that is uniform everywhere up to Δ*x* = *d*, after which it falls to zero (a *diffuse* kernel),
3. A decaying term, that decreases from a constant value at Δ*x* = 0 to zero at Δ*x* = *d* (a *decaying* kernel).

We also define *short-range* kernels, as any arbitrary function *h*(Δ*x*) that is nonzero only over scales|*x*| < *ϵ*_*i*_.

Any *simple* kernel *W*^*f*^ (Δ*x*) can be generally constructed as a linear combination of the above basic classes. In addition, *simple* kernels may also contain an added component of a *short-range* kernel.

To see that *simple* kernels will generally result in grid period ratios corresponding to Eq. (D1), we will examine the approximate Fourier transform structure for each component of the linear combination of *simple* kernels corresponding to a given length-scale *d*. We first demonstrate that each of the basic *simple* kernels will result in Fourier transforms that are sinusoidal functions with phase shifts and decaying envelopes and hence each basic *simple* kernel will satisfy Eq. (D1). We then show that short-range kernels present Fourier transforms that vary only at large scales, and can be ignored in our analyses of *simple* kernels. We then use these results to demonstrate that all *simple* kernels constructed as the above-described linear combination will have sinusoidal Fourier transforms and will satisfy Eq. (D1).

### 1 Localized kernels

For a general localized kernel *W*^*f*^ (Δ*x*) = *g*(|Δ*x*|− *d*) we obtain

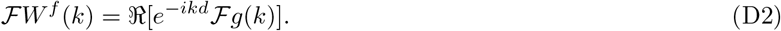

Since *g*(*x*) is supported over a scale *E*, the Fourier transform ℱ*g*(*k*) will only vary at scales *k* ∼ 1/*ϵ* ≫ 1/*d*. Thus for 1/*d* ≪ *k* ≪ 1/*ϵ*, we can approximate Eq. (D2) as

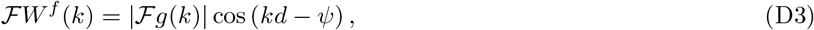

where *ψ* = arg[ℱ*g*(*k*)]. The local maxima of *FW*^*f*^ (*k*) will then occur at

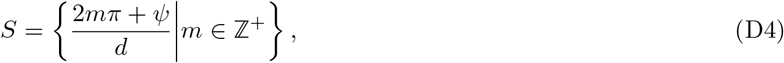

resulting in period ratios described by

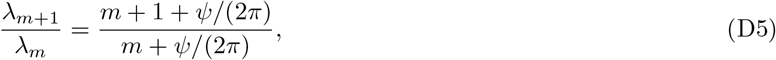

which is identical to Eq. (D1) for *ϕ* = *ψ*. We also note that we can now ascribe an interpretation to the phase angle *ϕ* — it is the phase difference between ℱ*W*^*f*^ (*k*) and cos(*kd*).

### 2. Diffuse kernels

We model a diffuse interaction kernel *W*^*f*^ (*n*_*DV*_) as

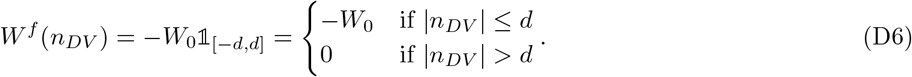

Corresponding to the discussion above, we look at the Fourier transform ℱ*W*^*f*^ (*k*)

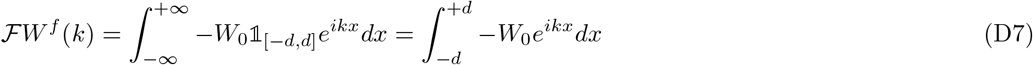

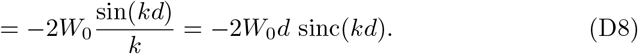

Note that once again, similar to Eqn. (B24), we obtain a functional form consisting of a periodic function (sin(*kd*)) that is multiplied by a decaying envelope 1/(*kd*). Ignoring the effects of the envelope function, the maxima of this function occur at

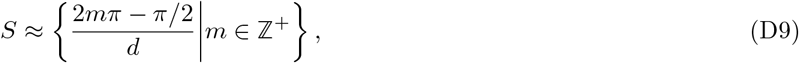

which immediately results in period ratios of the form

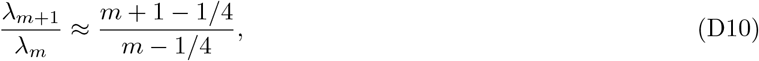

which corresponds to the result in Eq. (D1) for *ϕ* = *π*/2.

More precisely, the extrema of ℱ*W*^*f*^ (*k*) occur at *k*_*m*_*d* = *q* − 1/*q* − 2/3*q*^3^ + *O*(*q*^−5^) where 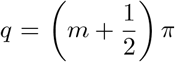 Notably, the errors decay approximately as 1/(*πm*), and thus for modules generated corresponding to *m* ≳: 2 will result in period ratios that approximate Eq. (D1) closely.

### 3. Decaying kernels

Decaying kernels with a scale *d* may be modeled as any monotonically decreasing function that decays from some constant *W*_0_ at Δ*x* = 0, to zero, at Δ*x* = *d*. For simplicity, we consider the simplest linear approximation to such a kernel, modeled as a triangular kernel. For additional subtleties in the treatment of other decaying kernels, see D 5 a The triangular kernel can be written as:

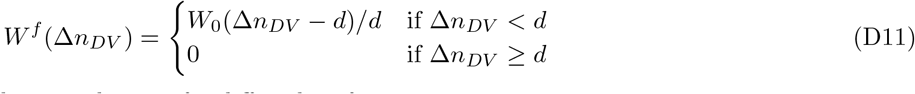

This function can be written as the convolution of 2 diffuse box functions:

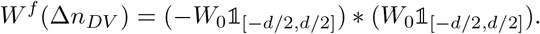

Thus, its Fourier transform is:

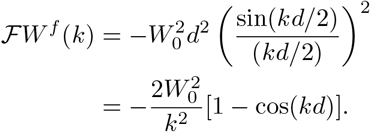

Once again, we obtain a simple trigonometric function, with maxima at

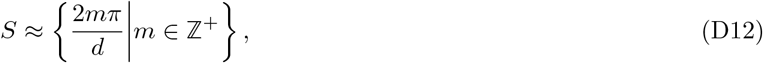

which immediately results in period ratios of the form

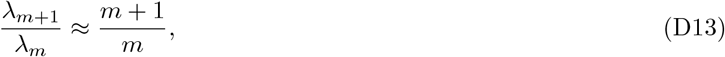

which corresponds to the result in Eq. (D1) for *ϕ* = 0.

### 4. Short-range kernels

For the case of a short-range kernel *W*^*f*^ (Δ*x*) that extends upto a scale *E*, we note from the Fourier uncertainty principle that the characteristic *k*-scales of ℱ*W*^*f*^ (*k*) will ∼1/*ϵ* ≫ 1/*d*. Thus, unlike the three other types of simple kernels discussed above, short range kernels do not have structure at the scale of 1/*d*. Since all relevant scales are much larger than 1/*d*, adding short range kernels to any of the other types of *simple* kernels will *not* change the structure of local maxima at scales of 1/*d*.

### 5. Arbitrary simple kernels

We now consider a general form for *simple* kernels, by constructing linear combinations of the above described three basic classes of *simple* kernels each corresponding to the same length scale *d* and additional short-range kernels.

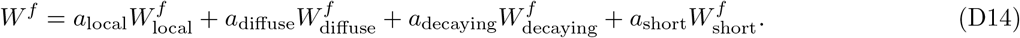

As demonstrated in the preceding sections, the Fourier transform *FW*^*f*^ (*k*) will be given as

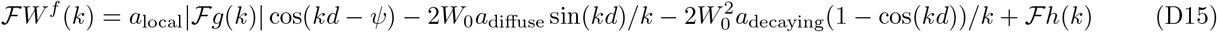

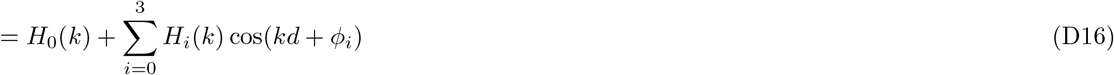

for some constants *ϕ*_*i*_, and some envelope functions *H*_*i*_(*k*) for *i* = 0, 1, 2, 3 that are slowly varying for *kd* ≳ 𝒪 (1). Under this approximation, ℱ*W*^*f*^ (*k*) is simply the sum of multiple sinusoidal waves with different phases and identical frequencies. Thus,

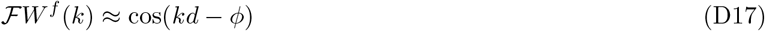

for some *ϕ* and *kd* ≳ *𝒪* (1). Hence, the maxima of ℱ*W*^*f*^ (*k*) occur at

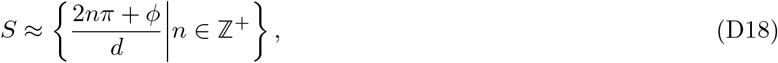

which immediately results in period ratios of the form Eq. (D1). Note that the approximations made above imply that there may be deviations from our results for the maxima corresponding to small *k* values — this may manifest as deviations in the largest period grid module away from Eq. D1.

#### a. Caveats

Clearly there exist *simple* kernels with Fourier trans forms that are not given by *FW*^*f*^ (*k*) ≈ cos(*kd*−*ϕ*). For example the Gaussian kernel, 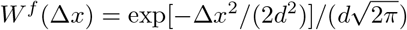 is a *simple* decaying kernel (since it has only a single scale *d*). Yet, its Fourier transform is simply ℱ*W*^*f*^ (*k*) = exp[−*k*^2^*d*^2^/2], which has only a single maximum! However, as we have shown earlier, there exist kernels that are arbitrarily close to the Gaussian kernel, whose Fourier transforms are given by powers of trigonometric functions, and hence have multiple regularly-spaced maxima with a spacing of ∼1/*d*. Similarly, there exist additional *simple* functions[118–120], ℱ*f* (Δ*x*), (like the Gaussian kernel) whose Fourier transforms *f* (*k*) have a small number of maxima. We hypothesize that for all such functions *f* (Δ*x*) there exist *simple* kernels *g*(Δ*x*) that are arbitrarily close to *f* (Δ*x*) and possess regularly spaced maxima.

### 6. Period ratios

Having demonstrated analytically that *simple* kernels result in a sequence of period ratios given by Eq. (D1), we now address the question of the mean period ratio over the sequence and over different values of *ϕ*. In the main text we have demonstrated that setting *ϕ* = 0 results in a detailed period ratio sequence that is in close agreement with the sequence of experimentally observed values. Here we consider the period ratios obtained for other values of *ϕ*, to demonstrate that the experimental observation of mean period ratios being approximated by 1.4 [24] emerges naturally from our setup.

From Eq. (D1), we obtained that the period ratio, *r*_*m*_ = *λ*_*m*+1_/*λ*_*m*_ can be written as

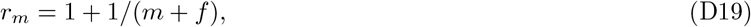

where *f* = *ϕ*/(2*π*). We ignore *m* = 1, since that results in a period ratio close to 2, which does not correspond to experimental observations. Averaging the period ratio over the next 4 modules (corresponding to *r*_*m*_ for *mϵ* {2 … 4}) results in

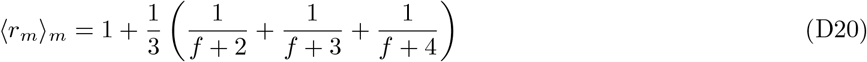

As can be seen in Fig. 15, this mean period ratio lies in the range [1.3,1.45], indicating that at all values of *ϕ*, the period ratio obtained from Eq. (D1) matches well with experimental observations. The average of these period ratios over all values of *ϕ* can also be calculated as

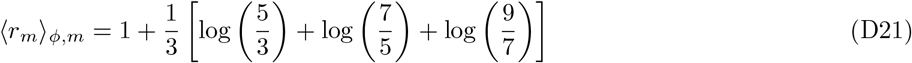

which is approximately equal to 1.37.

**FIG. 15.**
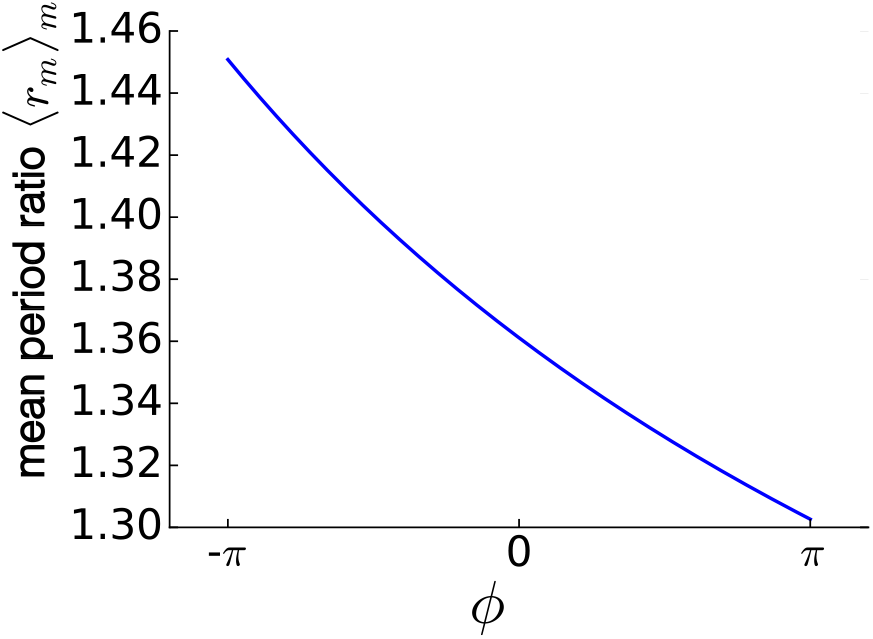
Mean grid-period ratios. Ratios of grid periods averaged over 4 modules as a function of the phase shift *ϕ* in Eq.(D1)

### 7. Module size; number of modules as a topological quantity

As discussed in the main text, peak-selection for modularization is a highly robust mechanism that is largely indifferent to system parameters such as the the particular forms of the fixed-scale interaction and the shape of the gradient. Here we provide an analysis of the number of modules, the scaling of module sizes, and the positions of module boundaries, which also exhibit the same robustness. Further, we also describe how this robustness may be interpreted as arising from a topological origin, similar to topological robustness in other physical systems like the quantum hall effect.

Recall that for the continuously graded kernel 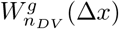 with characteristic spatial scale *σ*(*n*_*DV*_) at position *n*_*DV*_, the wave-vector of the formed pattern was proportional to 1/*σ*(*n*_*DV*_):

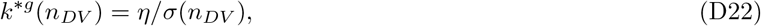

where *η* is an *n*_*DV*_ -independent constant that depends on only the particular form of the graded kernel. Let the spatial extent of the system be *n*_*DV ε*_ [0, *L*], with *σ*(*n*_*DV*_) monotonic such that *σ*_min_ = *σ*(0) ≤*σ*(*n*_*DV*_) ≤*σ*(*L*) = *σ*_max_.

We assume for simplicity that the fixed-scale lateral interaction is a *simple* kernel, such that ℱ*W*^*f*^ (*k*) ∼cos(*kd*− *ϕ*). Thus, the local maxima generated by *W*^*f*^ (*k*) occur at *k*_*n*_ (2*nπ* + *ϕ*)/*d*, where *n* are the natural numbers. As discussed in the main text, each of these local maxima is ‘selected’ in turn by the moving broad peak of the Fourier transform of the graded kernel, whose position according to Eq. D22 occurs at *k*^*\*g*^(*n*_*DV*_) = *η*/*σ*(*n*_*DV*_).

Notably, the selected maximum *k*_*m*_ will be robust to small perturbations in the selection function 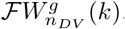 since *k*_*m*_ will remain quantized to one of the discrete values prespecified by the set {*k*_*n*_ ‖ *n ϵ* ℕ}. In this sense, the chosen maximum *k*_*m*_ (and hence the corresponding module) presents the hallmarks of a topologically protected state[1]. The topological number corresponding to a given module is the module number *m*, which is a topological invariant similar to a winding number[1][121].

The set of modules expressed through the length of the system corresponds to the set of local maxima *k*_*n*_ that lie within the range [*η*/*σ*_max_, *η*/*σ*_min_] that is delineated by the range of peak positions of the graded interaction. It follows that the maxima selected by the graded interaction obey:

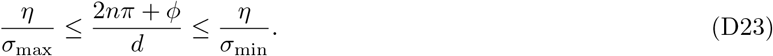

Thus, the set of formed modules are determined by the set of integers *n* that fit in the following interval:

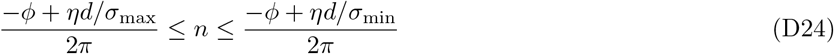

and hence the number of modules *N*_*mod*_ is:

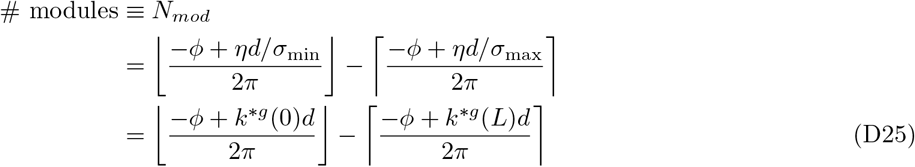

where ⌊ ⌋, ⌈ ⌉ indicate the floor and ceiling operations, respectively.

The above result leads to the following observations: First, the central quantity essential for determining the number of modules is the difference in the integer ratios of the fixed-scale interaction width to the extremal lateral interaction widths, *d*/*σ*_min_, *d*/*σ*_max_. Second, the number of modules depends only on the end-point values *σ*_min_, *σ*_max_ of the smoothly varying width *σ*(*n*_*DV*_) the graded interaction; notably, it does not depend on the detailed shape of *σ*(*n*_*DV*_). Moreover, if *σ*_min_, *σ*_max_ are varied smoothly (while *d* is held fixed), or if *d* is varied smoothly (while *σ*_min_, *σ*_max_ are held fixed), the number of modules will remain fixed, until the change becomes large enough to accommodate one additional or one less module. Thus, the number of modules is also a topological invariant of the system, through the module number *m*. Third, the number of modules does not depend on the system size *L*, or the number of neurons *n*_*DV*_ the system is discretized into (cf. Fig. 3f). Fourth, since the average module size will be *L*/*N*_*mod*_, the module sizes are extensive in *L*. Thus, for sufficiently large *L*, the module sizes can be orders of magnitude larger than the scales of the lateral interaction *d* and *σ*.

Note that the above argument on topological robustness of the modularization of the system is not restricted to the case of *simple* fixed-scale kernels. Indeed, for any fixed-scale interaction *W*^*f*^, the topological number *m* for any given expressed module will correspond to selecting the *m*^th^ maximum of ℱ*W*^*f*^ (*k*), for *k* > 0.

#### a. Module boundary locations

Following the peak-selection arguments made earlier, the module boundaries will occur at spatial locations that have *k*^*\*g*^(*n*_*DV*_) in between *k*_*n*_ and *k*_*n*+1_ (the specific location will depend on the particular forms of the kernels). As a zeroth order approximation, we can assume that the module boundaries will occur near (*k*_*n*_ + *k*_*n*+1_)/2,

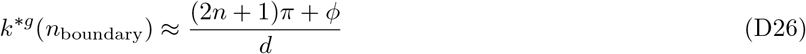

and thus

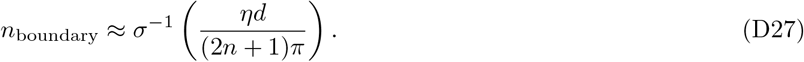

where *σ*^−1^ is the inverse function of *σ*(*n*_*DV*_), *σ*^*°*^*σ*(*x*) = *x*. Thus, while the specific positions of the module boundaries are dependent on the shape of the gradient *σ*(*n*_*DV*_), qualitative features such as the number of modules, module periods and module sizes are indifferent to the particular forms of the gradient (cf. Fig. 3f).

In (Fig. 4d), we vary the width of the *σ*(*x*) in two different ways: linearly along and in a square root along *n*_*DV*_. This leads to a shift in the module boundary locations that is predicted by fourier theory.

### 8 2D analysis

We have presented a majority of the above analysis for the case of one-dimensional grid cells. Here we briefly present the analogous computations for the Fourier transforms in two dimensions. We first demonstrate a classical result relating the Fourier transform of radially symmetric functions to the Hankel transform, which we shall then use to compute the relevant transforms. Consider the Fourier transform of a function *f* (**x**) = *f* (*x, y*)

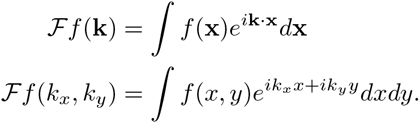

Define polar coordinates in real and Fourier space such that:

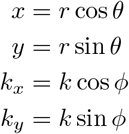

This leads to the dot product **k· x** to be simplified as

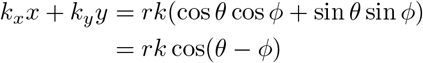

Thus,

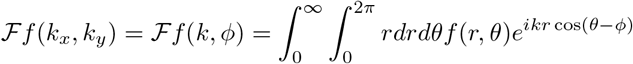

In all cases of interest, the function *f* is a kernel, and is hence a radially-symmetric real function *f* (*r, θ*) = *f* (*r*). Similarly, the Fourier transform ℱ*f* will also be a real radially-symmetric function ℱ*f* (*k, ϕ*) = ℱ*f* (*k*). Thus

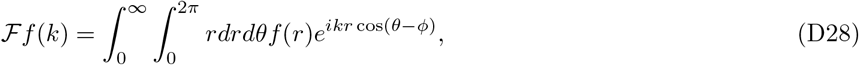

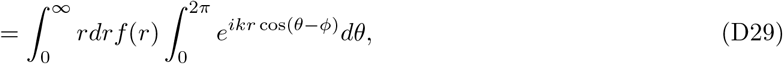

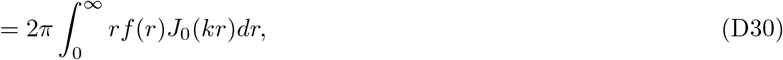

where *J*_0_ is the Bessel function of the first kind, defined by

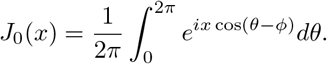

Equation (D30) defines the Hankel transform (of order zero) of *f* (*r*) — the radial component of the Fourier transform of the kernel *f* (**x**) is simply the Hankel transform of *f* (|**x**|).

For the localized gaussian secondary interaction, we can calculate the Fourier transform analytically.

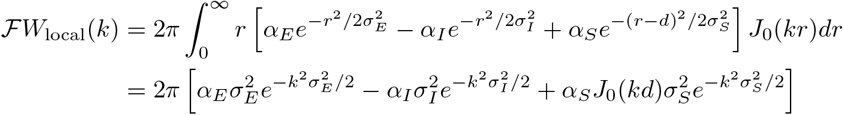

We can also analytically calculate the Fourier transform for a box-like interaction:

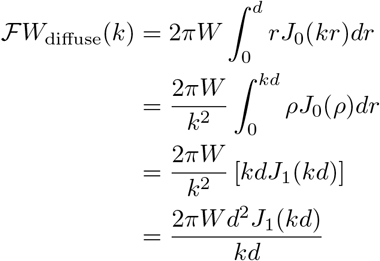

We can similarly also define a two-dimensional equivalent of the decaying kernel, as the convolution of the half-sized circular box kernel with itself. Thus, by applying convolution theorem to the result on diffuse kernels we obtain

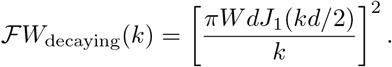

Note that *J*_0_(*x*) and *J*_1_(*x*) display qualitatively similar behavior to cos(*x*) and sin(*x*) respectively, apart from an amplitude modulation of the peaks — particularly, we note that the Bessel functions display approximately periodic maxima, which was the central property required for all of our results on modularization and peak selection to apply. We demonstrate this in Fig.16, where we show that the maxima of the Bessel functions are approximately periodic, and fit the form of Eq. (D1) well. In particular, note that the best-fit value of *ϕ* for *J*_0_(*k*) is approximately 0, which is similar to cos(*k*), and the best-fit value of *ϕ* for *J*_1_(*k*) is approximately *π*/4, which is similar to sin(*k*).

**FIG. 16.**
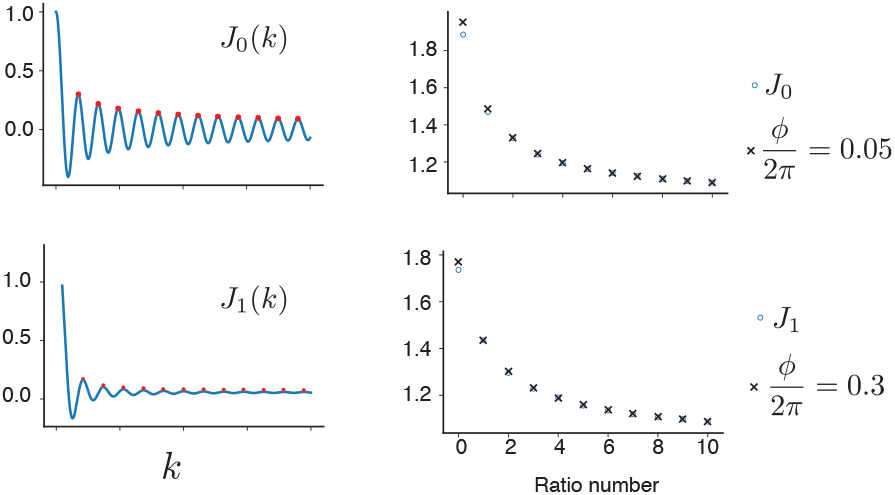
Bessel functions (left column) and period ratios for Bessel function maxima (right column) with their best-fit values of *ϕ* for the period ratios corresponding to Eq. (D1)

**FIG. 17.**
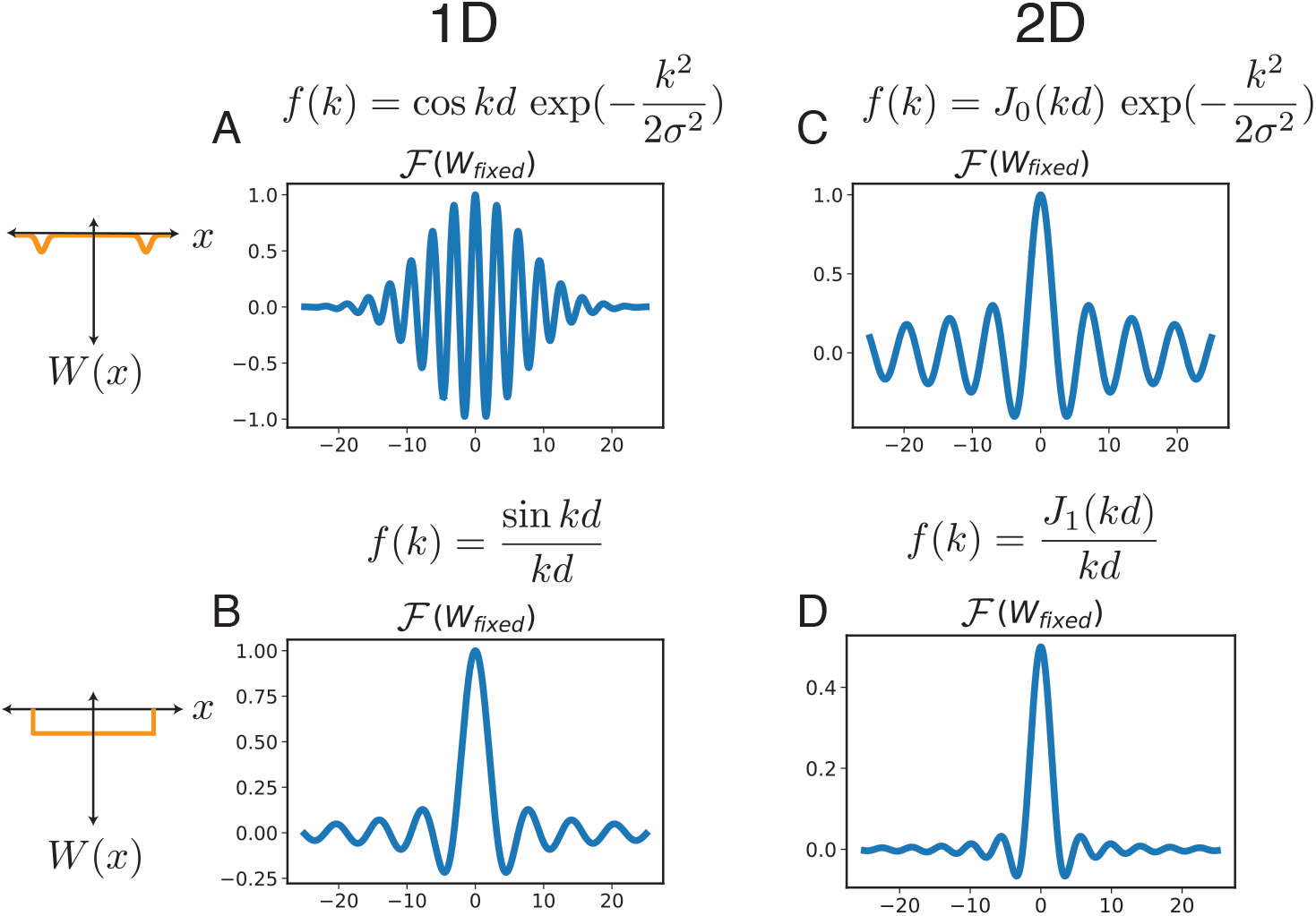
Fixed interactions(left, in orange) and their oscillatory Fourier transforms in 1D (left column) and 2D (right column).

We implemented a 2d simulation that generates 3 discrete modules as shown in Figure 18. For computational feasibility, the simulation was performed in 2 parts: one with 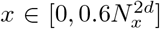 and the other with *x ϵ* 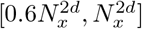 The weight matrices for each network were of size 100×1000 each. The weight matrix for a single large 100×2000 network would have contained 4*x*10^10^ elements, which we found prohibitively difficult and slow to run.

**FIG. 18.**
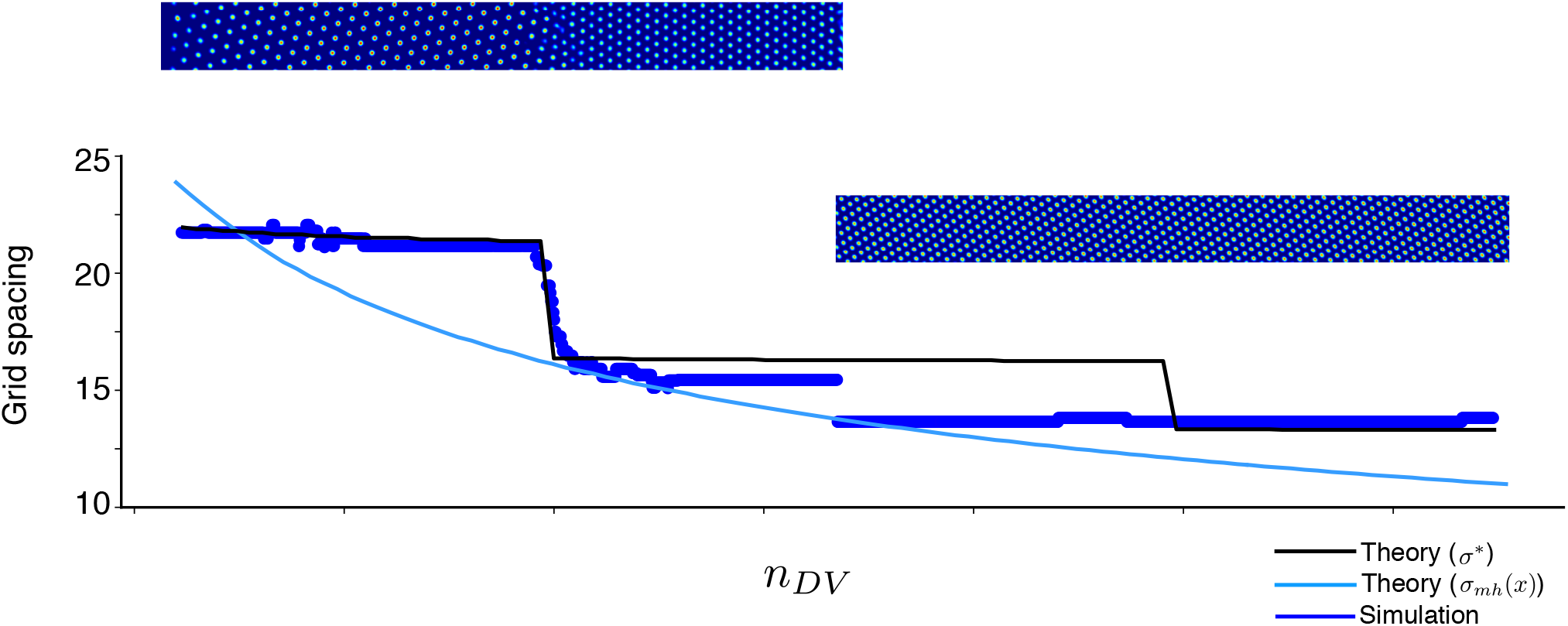
2d simulation with 3 modules: (top) Snapshots of population activity showing 3 discrete 2d grid modules, (bottom) plot of grid spacing and comparision with Hankel transform predictions. Grid spacing determined by calculating the (neural) spatial auto-correlation of the population firing activity.

Fig 19(a) shows another instance of a modular 2d network, the only difference being the value of *d*_*loc*_, which changed from 50 to 45. Fig 19(b) shows the same simulation with 2 distinct random initializations. The pair of resulting modules in each simulation have different relative orientations. Because finite size effects from our simulations also partially constrain the orientations of the modules (data not shown), we cannot make predictions about the relative orientations of the grid modules found in experiments [24].

**FIG. 19.**
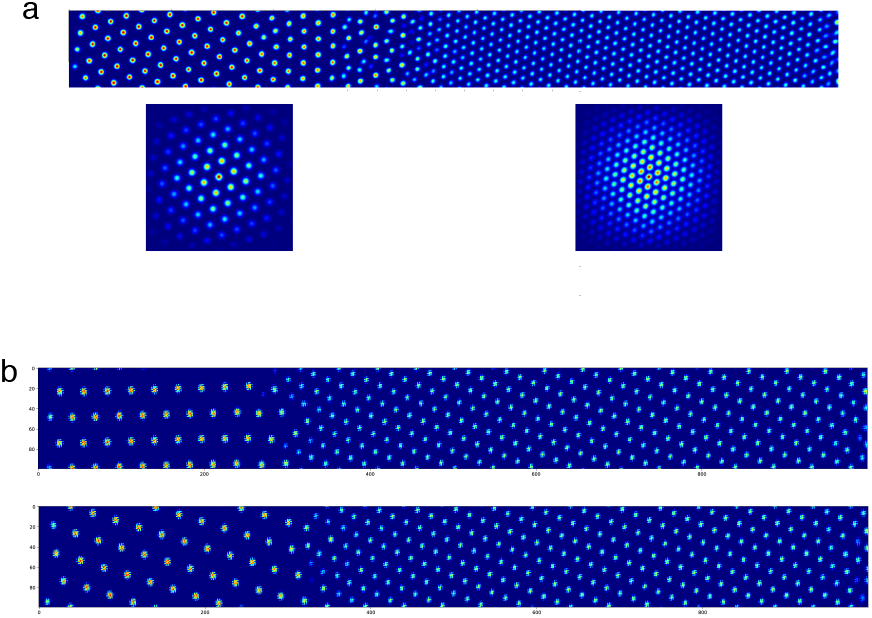
(a) Another instance of a spontaneously formed two dimensional network with parameters given in Table 4. (b) Two different random initializations of the network from Fig 2h show different relative orientations between the 2 formed modules.

### 9. Robustness to spatial noise

In the main text, we discussed how the topological robustness properties of peak selection result in the formed modules being stable to several forms of noise. Particularly, here we focus on the robustness to spatial heterogeneities in the lateral interaction kernels.

We first examine the robustness to spatial heterogeneities in the pattern forming kernel *W*^*g*^. To construct such an inhomogeneous pattern-forming interaction, we construct the noisy kernel at location **x**, by replacing the spatially homogeneous kernel *W*^*g*^[**x, x**′] = *W*^*g*^[**x** − **x**′], with a spatially heterogeneous kernel 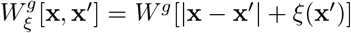 where *ξ*(**x**′) is a random number sampled independently for each spatial location **x**′ with mean zero and variance *ϵ*^2^. In Fig. 20d we present examples of such kernels for the case of *W*^*g*^[**x, x**′] described by the box function Eq. (13). Note how the independent sampling of *ξ*(**x**′) at each location results in a heterogenous kernel 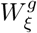 that varies in scale at different **x**, and is no longer radially symmetric.

**FIG. 20.**
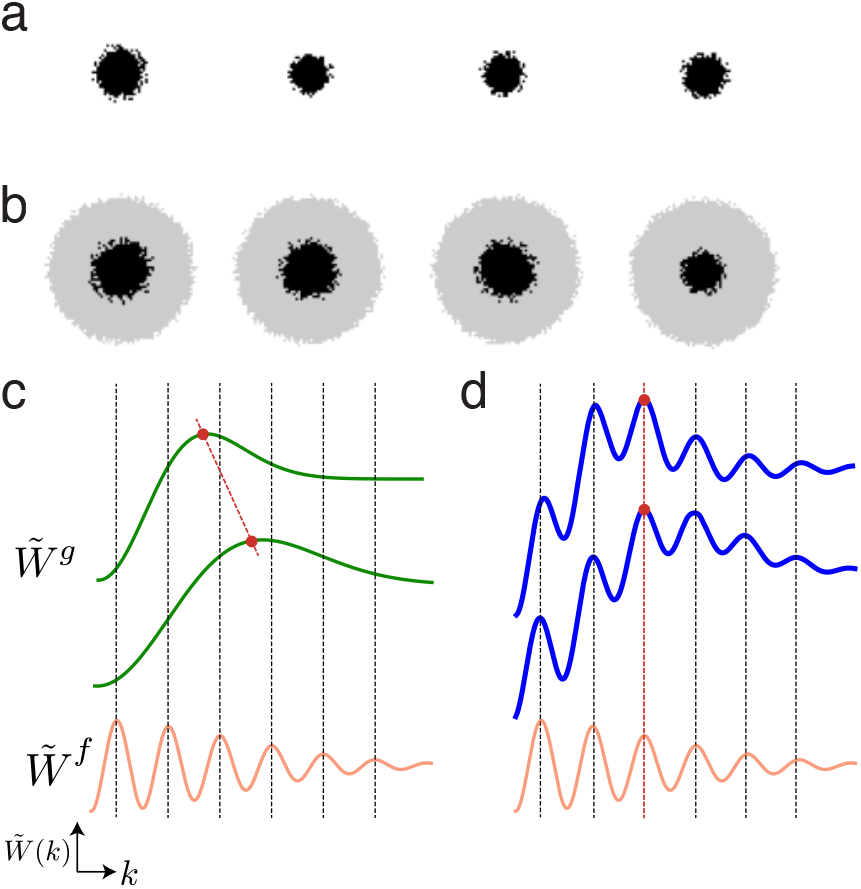
Noise robustness in peak selection process demonstrating how the additional of the smaller oscillatory fourier transform of the fixed interaction leads to no change in maxima despite smooth movement of the primary peak. (a) Example pattern forming interaction kernels from 4 neurons without a secondary fixed scale interaction (b) Example composite kernels from 4 neurons showing both the pattern forming interaction (black) and fixed scale interaction (grey)(c-d) The movement of the pattern forming interaction leads to a shift in the location of the gloabl maxima in the absence of a secondary interaction. This secondary interaction prevents any shift in the location of the global maxima when defined by the sum of the pattern forming interaction and the fixed scale interaction.

Recall that peak selection entails that the grid period at any location *n*_*DV*_ is dependent on the set of potential maxima defined by ℱ*W*^*f*^ (*k*), with a selection between these maxima performed by the broader peak of ℱ*W*^*g*^(*k*). If noise in the form of spatial heterogeneities are only introduced in ℱ*W*^*g*^ (and hence introduced in *W*^*g*^) this results in a noisy selection function. However, since the same maxima will be chosen for a range of selection functions (See Fig. 20a-b), the heterogeneity in *W*^*g*^ will not be manifested in the emergent grid period.

We next consider the addition of similar heterogeneities in the fixed-scale interaction as well, *W*^*f*^ (such as in Fig. 20c). Note that maxima induced by *simple W*^*f*^ are at *k*_*n*_≈ (2*nπ* + *ϕ*)/*d*, where *n* are the natural numbers, and hence the grid periodicity of the *n*^th^ module is given by *λ*_*n*_ ≈ *d*/(*n* + *ϕ*/2*π*). If we consider (*E*) noise added to *W*^*f*^ in the form of spatial heterogeneities, this would result in an (𝒪*ϵ*) error in the effective fixed-scale *d*. However, since *λ*_*n*_ is approximately *d*/*n*, thus the effective noise in periodicity of the *n*^th^ module, *λ*_*n*_, will be 𝒪 (*ϵ*/*n*). Thus, higher module numbers (corresponding to modules with smaller grid periods) have additional error correction beyond the robustness conferred by the topological nature of the peak selection process. This results in clean hexagonal firing fields despite inhomogeneities introduced in all lateral interactions as shown in Fig. 6.

### 10. Comparison of experimental observations with predicted period ratios

The general mechanism of peak-selection presented above describes how discrete modules can spontaneously arise in the presence of continuous gradients, by consideration of an additional fixed-scale lateral interaction *W*^*f*^. However, this mechanism does not provide any testable predictions for the ratio of grid periods unless additional assumptions are made. If indeed we assume that *W*^*f*^ is a *simple* kernel, i.e., *W*^*f*^ is primarily defined by a single spatial scale, then we demonstrated in SI Sec. D that the period ratios will be given by the simple formula, Eq. D1. In this section, we show that experimental observations of grid periods largely appear to match our predicted period ratios for *simple* kernels with *ϕ* = 0.

For verification of our main results on the predicted form of period ratios, we examine the literature for grid period measurements for multiple simultaneously measured grid modules in rats[24, 122–124]. We note that a large fraction of experimental observations of grid cells with more than one module measure only two modules. For a single pair of grid periods *λ*_1_ and *λ*_2_ > *λ*_1_, we can always explicitly solve for *ϕ* and *m* in Eq. (D1), to obtain

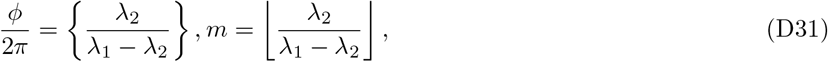

where {*x*} represents that fractional part of ⌊*x*⌋, and *x* = *x*−{*x*} represents the integer part of *x*. Thus, a single ratio, because it can always be fit by Eq. (D1), imposes no constraints on the accuracy of the expression.

It is possible to obtain a value of *ϕ* from Eq. (D1) and a single pair of periods; however, the estimate obtained from a single pair is not robust: *r*_*m*_ depends too sensitively on *ϕ*. For example, in [24], Rat 13388 exhibits grid periods of ≈53.24 cm and ≈43.00 cm (as estimated from SI Fig. 12b in [24]); Eq. (D1) then yields *ϕ*/(2*π*) = 0.199. Assuming a very small measurement error of ∼ 0.5*cm* in the larger period, such that if it were 53.75 cm instead of 53.24, would yield *ϕ* exactly equal to zero. A simple sensitivity analysis of the magnitude of error in estimating *ϕ* can be performed from Eq. (D31):

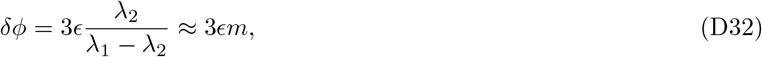

where *ϵ* represents the fractional error in the estimate of grid period. Thus, particularly for smaller grid periods (corresponding to larger *m*), even small errors in grid period estimation can result in a large error in *ϕ*, making the errorbars in the estimation of *ϕ* from a single pair of periods large.

To obtain results with significant statistical certainty, we focus our analysis on published experimental studies that measure at least 50 grid cells per animal, spanning at least 3 distinct modules. This restriction results in grid period data sets for three rats — we present kernel density estimates of the module periods for each of them in (Fig.21 (Fig. 21c corresponds to the data presented in the main text in Fig. 5).

**FIG. 21.**
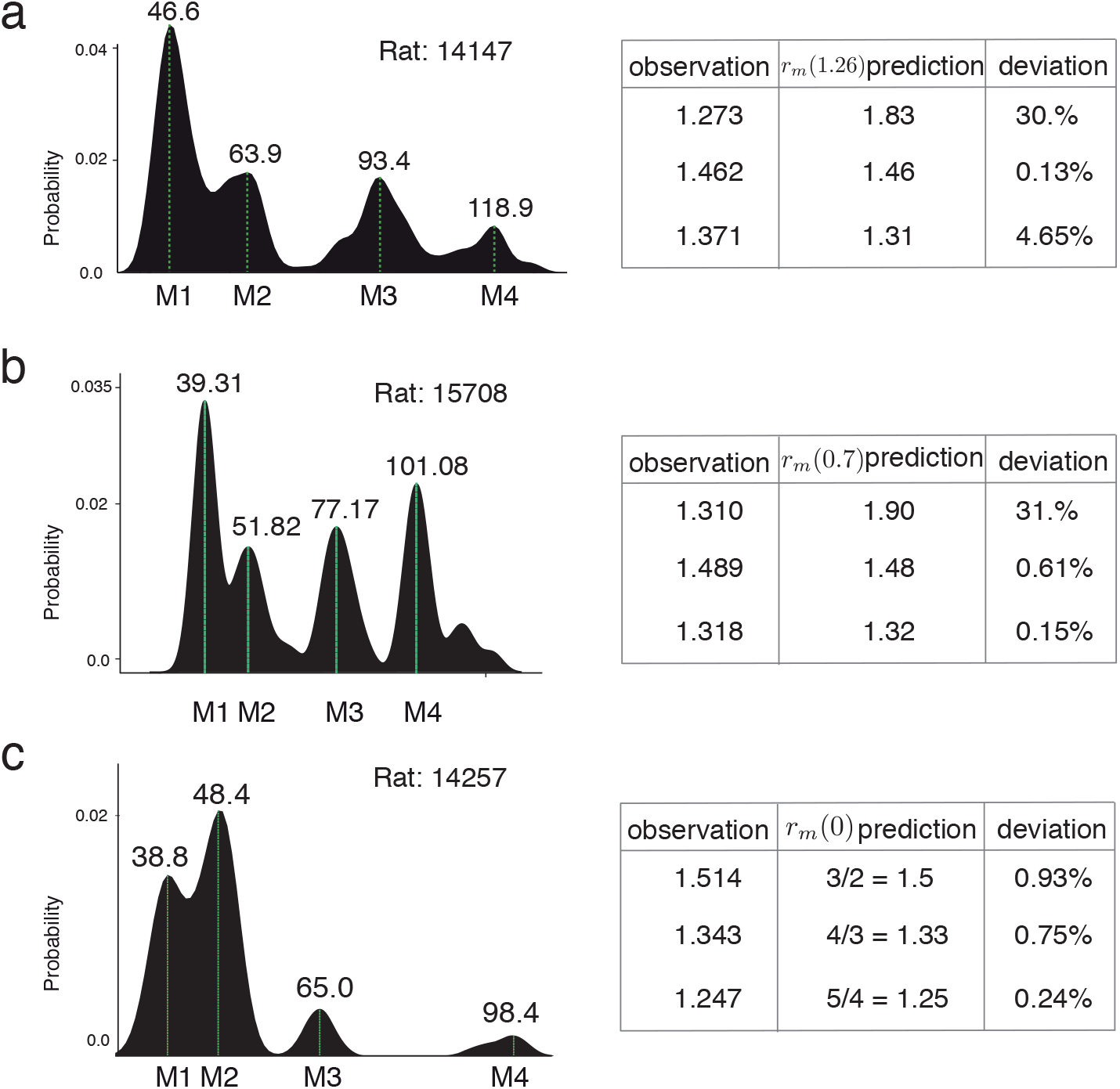
The 3 rats from Stensola *et al*. with 4 modules and their corresponding periods.

We have already demonstrated in Fig. 5 that Rat 14257 presents an extremely accurate match to the period ratio prediction for *ϕ* = 0 (i.e., predicted period ratios of 2, 3/2, 4/3, 5/4,…); in addition, Rat 14147 (observed period ratios of 1.27, 1.46 ≈3/2, 1.37≈ 4/3) and Rat 15708 (observed period ratios of 1.31, 1. ≈49 3/2, 1.32 ≈4/3) also match *ϕ* = 0 very well (*R*^2^ values of 0.999, 0.979, and 0.968 for Rats 14257, 15708, 14147 resp.) for all grid modules except for the module with the largest period.

Why is there an observed discrepancy for the grid module with the largest period? We propose four possible reasons for this discrepancy: Firstly, this discrepancy may be a result of the approximation made in arriving at Eq. (D17) — since the approximation is particularly accurate for *kd ≳*𝒪: (1), the potentialy mismatch would primarily affect only the largest grid period module. Secondly, as demonstrated in Sec. D9, the grid module corresponding to the largest grid period will have the least robustness to noise in the fixed-scale interaction, potentially introducing a large variance in the grid period for that module. Thirdly, as can be seen in Fig. 3h and Eq. (D32), the error in estimating the grid period for the first module (*m* = 1) is the most susceptible to errors in the value of *ϕ* Lastly, our predictions for grid period ratios Eq. (D1) are for the case of *simple* kernels that have a single spatial scale. A discrepancy at only the largest grid module may thus be suggestive of fixed-scale interactions that are primarily described by a single scale, with an additional low frequency perturbation at a larger spatial scale.

However, note that (particularly for Rats 14147 and 14257) there are relatively few grid cells observed from this largest period module, and the resulting uncertainty in period estimation may instead contribute to the error. In sum, apart from the possibility of some additional low frequency perturbations, the experimental data for rats with several simultaneously observed grid modules is largely consistent with the predicted period ratios for simple kernels with *ϕ* = 0.

Skipped modules: Sometimes, neural recordings can miss a module. This can cause a large deviation from our predictions. For example, for a set of 5 modules following period ratios M4/M5 = 1.20, M3/M4 = 1.25, M2/M3 = 1.33, M1/M2 = 1.5. If recordings had missed module M4, the measured ratios would be M1/M2 = 1.5, M2/M3 = 1.33, M3/M5 = 1.5.

However, we do note that available data on multiple modules with a statistically large number of grid cells per module are quite sparse. To obtain further verification of our theoretical results, including the prediction of Eq. (D1) and even more specifically the hypothesis that *ϕ* is close to zero, additional data with multiple simultaneously observed grid modules will be important.

## Appendix E Lyapunov Function

The energy function of continuous time neural networks can be written as [125]:

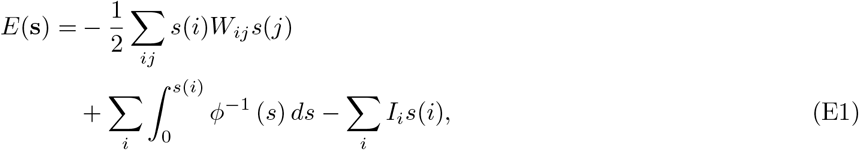

where **s** represents a vector of the synaptic activation at each neuron in the network, and *I*_*i*_ is the input bias to neuron *i*. For simplicity and since linear analysis does a remarkably good job in predicting the formed modules, let us restrict ourselves to the case of *ϕ* (*x*) = *x*. Also, since the system is locally translationally invariant, we know that the dominant modes are going to be periodic. Hence, we may evaluate the energy function of the network dynamics (in the linearized regime) by assessing the energy of the periodic neural activity modes:

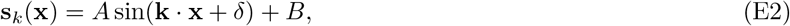

where 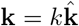 is an arbitrary Fourier space vector, and *A,B* and *δ* are arbitrary constants. For these modes, we can write the energy function in the continuum limit as:

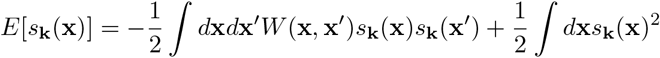

Assuming that the system size *L* is large,

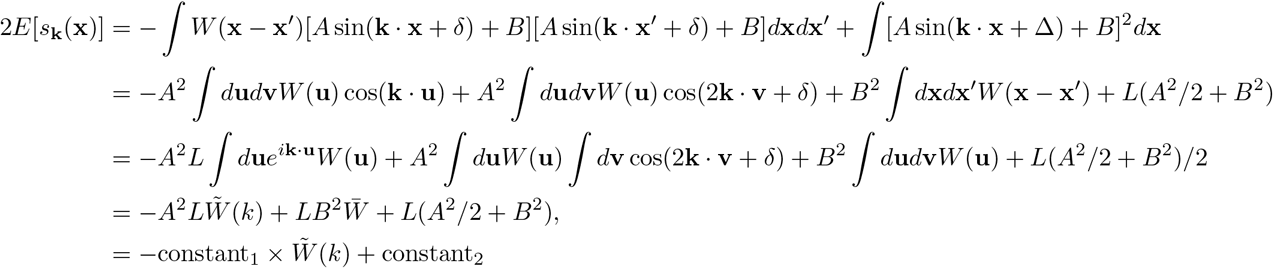

where have used the simple trigonometric identity, 2 sin(*C*) sin(*D*) = cos(*C* − *D*) − cos(*C* + *D*), and a change of variables, ∫ *d***x***d***x**′ = (1/2) ∫*d*(**x x**′)*d*(**x** + **x**) = *d***u***d***v**, with **u** = **x x**′ and 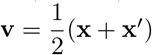

Thus, we obtain that the energy function *E*[*s*_**k**_] is a simple linear function of the Fourier transform 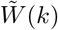 of the recurrent weight matrix. The minimum energy solution corresponds to the Fourier mode that maximizes 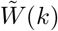. In other words, the dynamics is dominated by the *k*^*∗*^ that maximizes 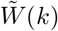. This result, derived from an energy landscape perspective, is equivalent to the result in Eq. (B10), which we obtained earlier via perturbation analysis.

## Appendix F General formulation of module formation dynamics: Discrete peak selection via loss minimization

In Sec. E, we demonstrated how the pattern formation on the neural sheet can be derived via an energy minimization approach. Here, we use an energy landscape view to describe how loss function minimization results in modular solutions.

The key components for spatially modular solutions to arise from energy minimization are as follows: 1) A spatially-independent loss function *f* (*θ*) with multiple local maxima and minima; 2) A gradient in a spatially-dependent variable, *θ*_0_(*x*); and 3) A coupling between the system parameters *θ* and *θ*_0_, that results in a combined loss function

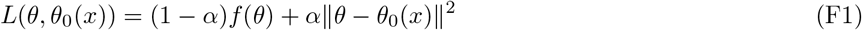

Under appropriate constraints on *f* (*θ*), solving the following optimization at each *x*

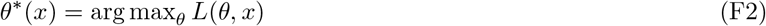

will produce discrete, step-like changes as a function of *x*. This happens because the smooth minimum given by the | |*θ* − *θ*_0_(*x*) | | ^2^ term effectively selects one of the local minima in *f* (*θ*) as the global minimum. As the function | |*θ* − *θ*_0_(*x*) | | ^2^ slides smoothly along with *x*, the peak of *f* (*θ*) selected as the global minimum remains the same for some time, then jumps abruptly. These step-like changes are modular solutions to the global optimization problem. The energy function defined in Eq. (F1) can be viewed as a regularized optimization problem, with the spatially-dependent regularizer | |*θ* − *θ*_0_(*x*) | | ^2^ acting as a prior that selects one of the minima of *f* (*θ*) at each location (Fig. 22).

**FIG. 22.**
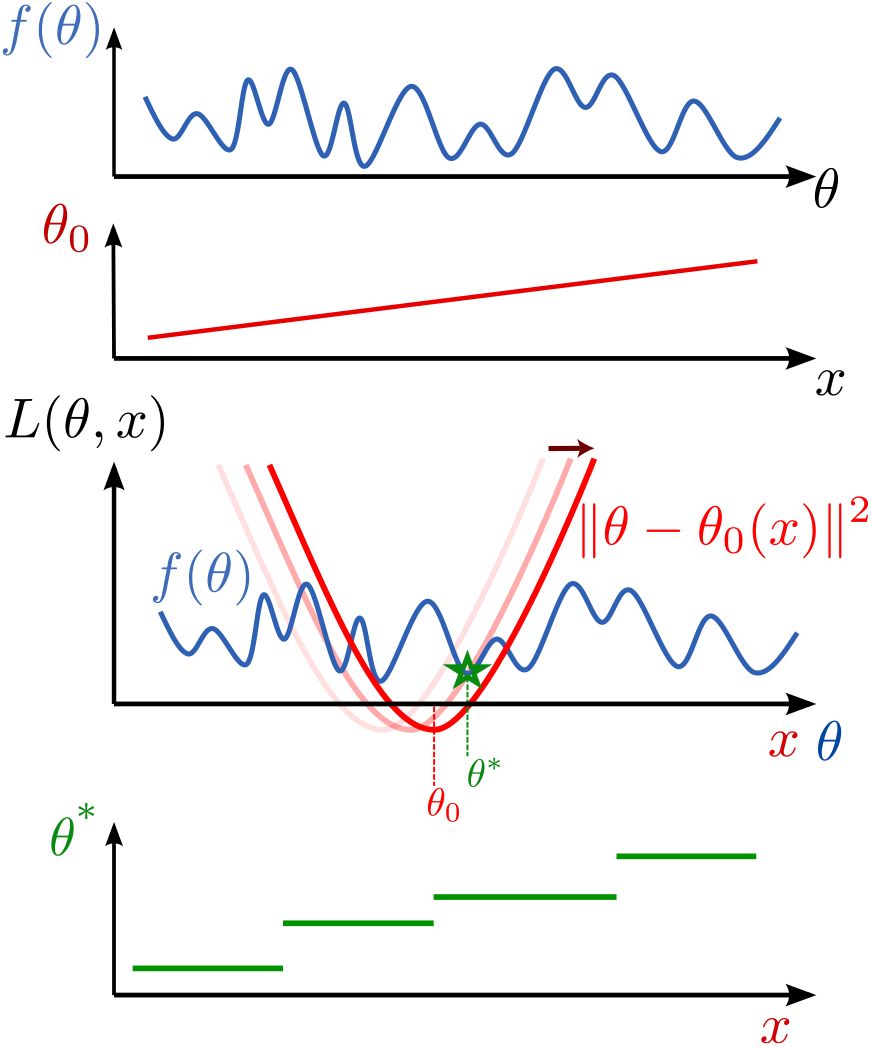
A general setting for peak-selection. Assuming a loss function *f* (*θ*) (blue) and a spatially dependent quantity *θ*_0_ (red), a combined loss function *L*(*θ, x*) can be constructed such that the *x*-dependent optimizer of *L*(*θ, x*) will be modular (green), since it will be constrained to correspond to one of the minima of *f* (*θ*).

The correspondence of this general picture with the peak selection mechanism described in the main text follows directly with the following identifications: the spatially independent nonlinear loss function *f* (*θ*) with the fixed-scale interaction *W*^*f*^ ; the spatially varying parameter prior *θ*_0_(*x*) with the graded scale *σ*(*n*_*DV*_) of the pattern-forming kernel; the combined loss *L*(*θ, x*) with the full kernel *W*_*nDV*_ ; and the spatially-varying, multi-step-like set of optima *θ*^*∗*^(*x*) with the grid periods *λ*^*∗*^(*x*), respectively. Similar to peak selection for grid cells, the formed modules in this generalized setting will also inherit topological robustness and stability.

We demonstrate a numerical example of this in Fig. 7, where we construct *f* (*θ*) as a random sample from a Gaussian process with a radial basis function kernel, and simulate gradient descent dynamics on the loss function *L*(*θ, θ*_0_(*x*)). To prevent the dynamics from getting stuck in local minima of *L*, we simulate the gradient descent first purely on the regularization term, with gradually increasing strength of the rugged loss function, through gradually decreasing *α* with increasing time.

Although we primarily focused on the peak selection process in Fourier space for multi-periodic patterning in grid cells, we also showed that it has a general formulation in terms of dynamics on an energy landscape: One (spatially invariant) interaction sets up an optimization problem with multiple local minima, while a second (spatially graded) interaction defines a locally shallow single-optimum landscape, with a smoothly shifting optimum as a function of space. Thus, the shallow optimum selects one of the narrow local optima as the global optimum, with discontinuous jumps to the next local minimum even as the parameters vary smoothly. This analytical formulation provides a simplifying mathematical perspective on how smooth gradients could lead to discrete patterning and modular specialization in the brain and body [31, 40, 62].

## Appendix G The emergence of modules corresponds to the formation of localized eigenvectors

As has been observed before [126], a neural network endowed with slowly varying local interactions shows diverse timescales that are spatially localized: different parts of the network respond with disparate temporal dynamics. We also find a localization of eigenvectors in our multi-module grid network, Fig. 23A. Similar to [126], our interaction matrix has a locally circulant form (due to the slowly varying gradient in lateral inhibition width). This is a signature of a phase transition, similar to the Anderson localization transition in condensed matter physics [53]. The eigenvectors for a regular pattern forming interaction in traditional continuous attractor models are delocalized fourier waves which are then transformed into localized fixed-wavelength gaussian wavepackets with the addition of the gradient and fixed scale interaction.

**FIG. 23.**
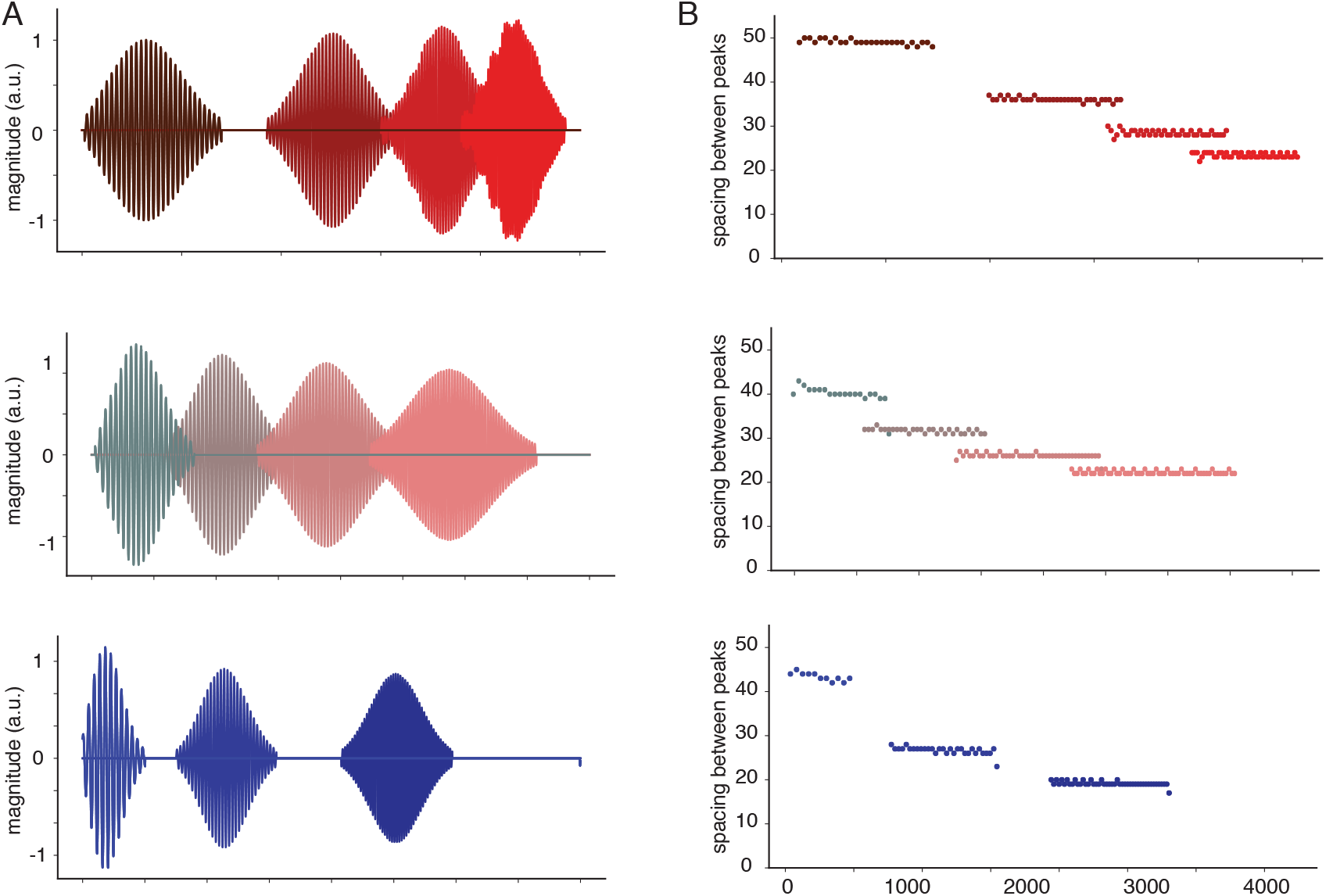
Localization of eigenvectors: A) Eigenvectors of various one-dimensional interaction weight matrices along with the corresponding inter-peak spacings are localized, B) The periodicity within an eigenvector is constant

We find that in the resulting set of localized eigenvectors, each has a different but constant period, Fig. 23B. These periods exactly match the spatial periods of the modules formed in steady state. In sum, the locally circulant matrix gives rise to eigenvector localization, and the localized eigenvectors correspond to the modules.

